# Inhibiting xCT/SLC7A11 induces ferroptosis of myofibroblastic hepatic stellate cells and protects against liver fibrosis

**DOI:** 10.1101/2019.12.23.886259

**Authors:** Kuo Du, Seh Hoon Oh, Tianai Sun, Wen-Hsuan Yang, Jen-Tsan Ashley Chi, Anna Mae Diehl

## Abstract

**Background and Aims:** Liver fibrosis develops in the context of excessive oxidative stress, cell death and accumulation of myofibroblasts (MFs) derived from hepatic stellate cells (HSCs). Ferroptosis is a type of regulated cell death that can be caused by inhibiting the cystine/glutamate antiporter xCT. However, while xCT is induced in various liver diseases, its role in HSC activation and liver fibrosis is unknown. We hypothesized that xCT is required for HSCs to antagonize ferroptosis and remain myofibroblastic.

**Methods:** xCT activity was disrupted by siRNA or pharmacological inhibitors in MF-HSC cell lines to determine its effect on redox homeostasis, growth, myofibroblastic activity and viability. xCT expression was then determined by RNA sequencing and RT-PCR during primary HSC activation, and its role in HSC trans-differentiation was assessed. For comparison, xCT expression and function were also determined in primary hepatocytes. Finally, the roles of xCT in HSC accumulation and liver fibrosis were assessed in mice treated acutely with CCl4.

**Results:** Inhibiting xCT in MF-HSCs decreased intracellular glutathione (GSH), suppressed growth and fibrogenesis, and induced cell death. These effects were rescued by antioxidants, an iron chelator, and a canonical ferroptosis inhibitor, but not by inhibitors of apoptosis or necrosis. xCT was dramatically up-regulated during primary HSC activation, and inhibiting xCT suppressed myofibroblastic trans-differentiation and induced ferroptosis. In contrast, healthy hepatocytes were relatively insensitive to ferroptosis induced by xCT inhibition. In vivo, inhibiting xCT systemically reduced MF-HSC accumulation and liver fibrosis after a single dose of CCl4 without exacerbating liver injury or reducing hepatocyte regeneration.

**Conclusion:** Compared to healthy hepatocytes, MF-HSCs are exquisitely sensitive to ferroptosis induced by inhibiting xCT. In acutely injured livers, systemic inhibitors of xCT can inhibit fibrosis without worsening liver injury. Further research is needed to determine if this therapeutic window remains sufficiently robust to safely target MF-HSCs and inhibit fibrogenesis in chronically injured liver.

## Introduction

The liver, more than any other organ, has an incredible capacity to regenerate after injury. However, regenerative responses can be accompanied by a futile wound-healing process in chronic liver diseases, leading to excessive accumulation of extracellular matrix, i.e., fibrosis. Progressive liver fibrosis results in cirrhosis and thus, it is the final common pathway of chronic liver diseases of various etiologies, causing over 1.4 million death each year.^1, 2^ Myofibroblasts that secrete large amount of extracellular matrix proteins are the drivers of liver fibrosis. Hepatic stellate cells (HSCs) are the dominant source of these fibrogenic myofibroblasts.^3^ HSCs normally exist in a quiescent, non-proliferative state. However, when the liver is injured, HSCs trans-differentiate into proliferative, migratory and fibrogenic myofibroblastic hepatic stellate cells (MF-HSCs). Therefore, the mechanisms that control accumulation of MF-HSCs are attractive therapeutic targets for liver fibrosis.^3^ Previously, we demonstrated that the myofibroblast transition requires metabolic alterations that enable HSCs to catabolize glutamine to generate glutamate^4^.

Ferroptosis is a recently identified form of regulated cell death that is characterized by disturbed iron and redox homeostasis and increased lipid-peroxidation.^5^ It is distinct from other forms of cell death morphologically, biochemically and genetically, and has been implicated in both physiological and pathological processes, including neurodegenerative diseases, kidney injury and cancers.^6^ Studies of ferroptosis have identified some key regulators of the process by characterizing the mechanism of action of ferroptosis inducers such as erastin and RSL3.^5^ Erastin initiates ferroptosis by inhibiting xCT (encoded by *SLC7A11*), a cell surface amino acid antiporter that exports intracellular glutamate in exchange for extracellular cystine.^6^ Reducing cystine uptake decreases its intracellular conversion to cysteine, the rate-limiting amino acid for biosynthesis of GSH, leading to impaired function of glutathione peroxidase 4 (GPX4), and subsequent propagation of lipid peroxidation mediated by free active iron.^6^ RSL3 acts downstream of xCT to induce ferroptosis by directly inhibiting GPX4.^6^ Although xCT was reported to increase during toxic liver injury and in hepatocellular carcinoma (HCC),^7–9^ its role in MF-HSC has never been investigated and thus, the impact of xCT activity on liver fibrosis progression remains unclear.

Therefore, in the current study, we investigated the roles of both xCT and ferroptosis in HSC trans-differentiation and liver fibrosis. By deploying an array of complementary tools (human and rat MF-HSC cells lines, primary mouse HSCs and hepatocytes, and a mouse model of acute CCl4-induced liver fibrosis), we demonstrated that xCT is induced during the myofibroblastic differentiation process and required for MF-HSCs to maintain their myofibroblastic gene expression signatures. We found that MF-HSCs require xCT to resist ferroptosis in culture, while cultured primary hepatocytes are resistant to ferroptosis induced by xCT inhibitors. Furthermore, we showed that similar differential sensitivity to xCT inhibition occurs *in vivo* and thus, might be exploited to inhibit liver fibrosis in mice.

## Methods and Materials

### Cell culture studies

Primary HSCs were isolated from healthy 12-16 week old C57BL/6 mice. HSCs were cultured for up to 7 days in Gibco® DMEM containing 10% fetal bovine serum, and 1% penicillin-streptomycin (Life Technologies, Grand Island, NY) to induce activation. To test the role of xCT in HSC activation, cells were treated with the xCT inhibitor erastin (5μM) or its vehicle DMSO (0.1%) from day 4 for 3 days. To test whether quiescent HSCs are sensitive to xCT inhibition, cells were treated with the xCT inhibitor erastin (5μM) or its vehicle DMSO (0.1%) immediately after fresh isolation for 3 days. To test the role of xCT, glutaminase 1 (GLS1) and ferroptosis in MF-HSC cell lines, human MF-HSCs (LX2 cells from Scott Friedman, Mount Sinai School of Medicine) and rat MF-HSCs (8B cells from Marcos Rojkind, George Washington University, Washington, DC) were treated with 20nM nontargeting siRNA or GLS1 siRNA or xCT siRNA (Dharmacon), or 0.5 μM GPX4 inhibitor 1S, 3R-RSL3, or xCT pharmacological inhibitors (e.g. 1-10 μM erastin, 0.25-1 mM SASP), or GLS1 inhibitor (1μM CB839 or 10 μM BPTES) or the vehicle control DMSO (0.1%).

To further validate the role of xCT and ferroptosis in HSCs, in some of the above experiments the culture medium was supplemented with antioxidant NAC (5 mM) or β-mercaptoethanol (50 μM), or with inhibitors of ferroptosis (ferrostatin, 2 μM or the iron chelator deferoxamine,10 μM), or with the apoptosis inhibitor Z-VAD-FMK (10 μM) or the necroptosis inhibitor Necrostain-1 (0.5 μM). To compare the ferroptosis sensitivity of primary hepatocytes to HSCs, primary mouse hepatocytes were isolated from adult healthy mice. Cells were treated with the direct xCT inhibitor erastin (5μM; 10μM), the direct GPX4 inhibitor RSL3 (0.5 μM) or vehicle DMSO (0.1%) for 3 days. Cell death was assessed by CellTox™ Green Cytotoxicity Assay (Promega) according to manufacturer’s instructions. Cell growth/viability was monitored with Cell Counting Kit-8 according to manufacturer’s instructions (CCK8, Dojindo Molecular Technologies). For RT-PCR and western blot assay, cell medium was first removed and the cells were washed with PBS for 2 times to clear the cell debris. Only attached surviving cells were harvested.

### GSH and glutamate measurements

The intracellular GSH level was determined using the GSH/GSSG-Glo™ Assay kit (Promega, V6611), and the intracellular glutamate level was determined using the Glutamine/Glutamate-Glo™ Assay kit (Promega, J8022).

### Lipid peroxidation assay using flow cytometry

Lipid ROS levels were determined by C11-BODIPY assay according to the manufacturer’s instructions (D3861, ThermoFisher Scientific). Briefly, cells were treated with xCT inhibitors (1-2 μM erastin; 0.25-0.5 mM SASP) or the vehicle control DMSO (0.1%) for 24-48h. The medium was then replaced with 10 mM C11-BODIPY-containing medium for 1 h. Next, the cells were harvested by trypsin and resuspended in PBS plus 1% BSA. Lipid ROS level was examined by flow cytometry analysis (FACSCantoTM II, BD Biosciences).

### Quantitative real time PCR (qRT-PCR)

Total RNA was isolated from whole liver tissue or cultured cells using Trizol reagent. After determining the concentration and quality of the RNA, complementary DNAs (cDNA) were generated using Superscript II Reverse Transcriptase (Life Technologies) according to the manufacturer’s instructions. Each cDNA sample was assayed by qRT-PCR using SYBR Green Super-mix using primers listed in **Supplementary Table 1**. Results were normalized to the housekeeping gene S9 based on the threshold cycle (C*t*) and relative fold change was determined using the 2^-ΔΔCt^ method.

### Immunoblot analysis

Protein was extracted from isolated cell pellets or whole liver tissue using RIPA buffer with protease inhibitors (Sigma-Aldrich). Equal amounts of protein were loaded and separated by SDS-PAGE gel electrophoresis on 4%-20% Criterion gels (BioRad, Hercules, CA), and then transferred to PVDF membranes, and incubated with the following primary antibodies: αSMA (Abcam, ab32575), Vimentin (Abcam, 92547), Collagen I (Cell signaling, 84336), xCT (Cell signaling, 12691; or Invitrogen, PA1-16893), caspase-3 (Cell signaling, 9662); β-tubulin (Abcam, ab6046) or GAPDH (Santa Cruz, sc-47724) overnight at 4°C. Blots were visualized with HRP-conjugated secondary antibodies.

### Histopathological analysis, Immunohistochemistry (IHC) and Immunocytochemistry (ICC)

Liver tissue was fixed in formalin, embedded in paraffin, and cut into sections. The slides were stained with H&E and evaluated for hepatocyte injury. Liver fibrosis was assessed by Picrosirius red (#365548, Sigma) staining according to the manufacturer’s suggestions.

For IHC, slides were dewaxed, hydrated, and incubated for 10 min in 3% hydrogen peroxide to block endogenous peroxidase. Antigen retrieval was performed by heating in 10 mmol/L sodium citrate buffer (pH 6.0) for 10 min. Sections were blocked in Dako protein block solution (X090, Agilent Technologies, Santa Clara, CA) for 1hr and incubated OVN at 4°C with indicated primary antibodies: αSMA (Dako, M0851); Desmin (Abcam, ab15200); CyclinD1 (Abcam, ab134175). Polymer-horseradish peroxidase secondary antibodies were applied for 1h at room temperature, and Dako *3*,*3*′-*Diaminobenzidine* Substrate Chromogen System was used for detection. Staining was quantified by morphometry (MetaView software, Universal Imaging Corp) using 5 randomly chosen fields at ×4 magnification or 10 randomly chosen fields at ×10 magnification per section for each mouse.

For ICC, cells were washed, fixed in 4% paraformaldehyde, permeabilized, blocked with normal goat serum, and incubated overnight with antibodies to αSMA (M0851) or Col1α1(ab34710) or xCT (Invitrogen, PA1-16893). Cells were washed in PBS and incubated with Texas Red goat anti-mouse IgG or Alexa Fluor 488 goat anti-rabbit IgG (H+L, ThermoFisher Scientific) for 1h at RT. 4’,6-diamidino-2-phenylindole (*DAPI*) was used to visualize nuclei. Images were acquired and processed using a Zeiss LSM710 inverted confocal microscope system.

### Animal studies

To acutely induce liver fibrosis, C57BL/6 mice (Jackson Laboratories, Bar Harbor, ME) were injected intraperitoneally with corn oil or 1200 mg/kg CCl4 (n=4 mice/group). To determine if inhibiting xCT altered liver injury and MF accumulation, erastin (30 mg/kg) or its vehicle (10%DMSO in PBS) was intraperitoneally administered at 9h and 30h post-CCl4 to avoid a potential effect on CCl4 toxic metabolism. The dose and route of administration of erastin were based on previous studies using erastin *in vivo.*^10–12^ Mice were sacrificed 48h after treatment with vehicle or CCl4 (n=5 mice/group). At sacrifice, blood was obtained for subsequent measurement of AST and ALT; HSCs and liver tissue were also harvested: HSC were immediately analyzed for expression of xCT or MF markers or placed into culture; liver tissues were fixed in phosphate-buffered formalin for histological analysis, or flash-frozen in liquid nitrogen and stored at −80 °C until analysis. All studies were approved by the Duke University Institutional Animal Care and fulfilled National Institutes for Health and Duke University IACUC requirements for humane animal care.

### Statistics

Data were expressed as mean ± SEM. Statistical significance between two groups was evaluated using the student’s t test, while comparisons of multiple groups were assessed by one-way analysis of variance (ANOVA), followed by Student–Newman–Keul’s test. *p* ≤ 0.05 was considered to be statistically significant.

## Results

### xCT is critically required for MF-HSC myofibroblastic activity

xCT, encoded by *SLC7A11*, is the plasma membrane antiporter that imports cystine while exporting glutamate. xCT plays critical roles in GSH biosynthesis, ROS defense, glucose and glutamine consumption, mitochondrial respiration, lipid metabolism, metabolic reprogramming and cell growth in various cancer cell lines and animal models of cancer.^13^ Like cancer cells, other types of highly proliferating cells express xCT, and xCT^-/-^ fibroblasts failed to survive under conventional culture conditions due to an insufficient uptake of cystine.^14^ While quiescent HSCs are known to become proliferative during their trans-differentiation into fibrogenic MFs, the role of xCT in MF-HSCs remains virtually unknown. We hypothesized that xCT is critically required for MF-HSC to maintain their myofibroblastic activity. To test this hypothesis, human MF-HSCs (LX2 cells) were treated with *SLC7A11* siRNA for 5 days to knockdown xCT expression. Measurement of xCT (*SLC7A11*) mRNA (**Fig.1A**) and protein expression (**Fig. 1B**) demonstrated the knockdown efficiency of the siRNA. As expected, xCT knockdown significantly elevated intracellular glutamate levels (**Fig.1C)** but reduced intracellular GSH (**Fig.1D**). Most interestingly, xCT depletion also reduced the growth of MF-HSC (**Fig.1E; Suppl. Fig. 1A**). Furthermore, xCT depletion also decreased expression of the MF-HSC markers alpha smooth muscle actin (αSMA, encoded by ACTA2) and vimentin (VIM) **(Figs. 1F, G),** but increased the expression of quiescent markers peroxisome proliferator receptor-gamma (PPARγ) and E-cadherin (E-cad) **(Suppl. Fig. 1B)**.

**Figure 1.**
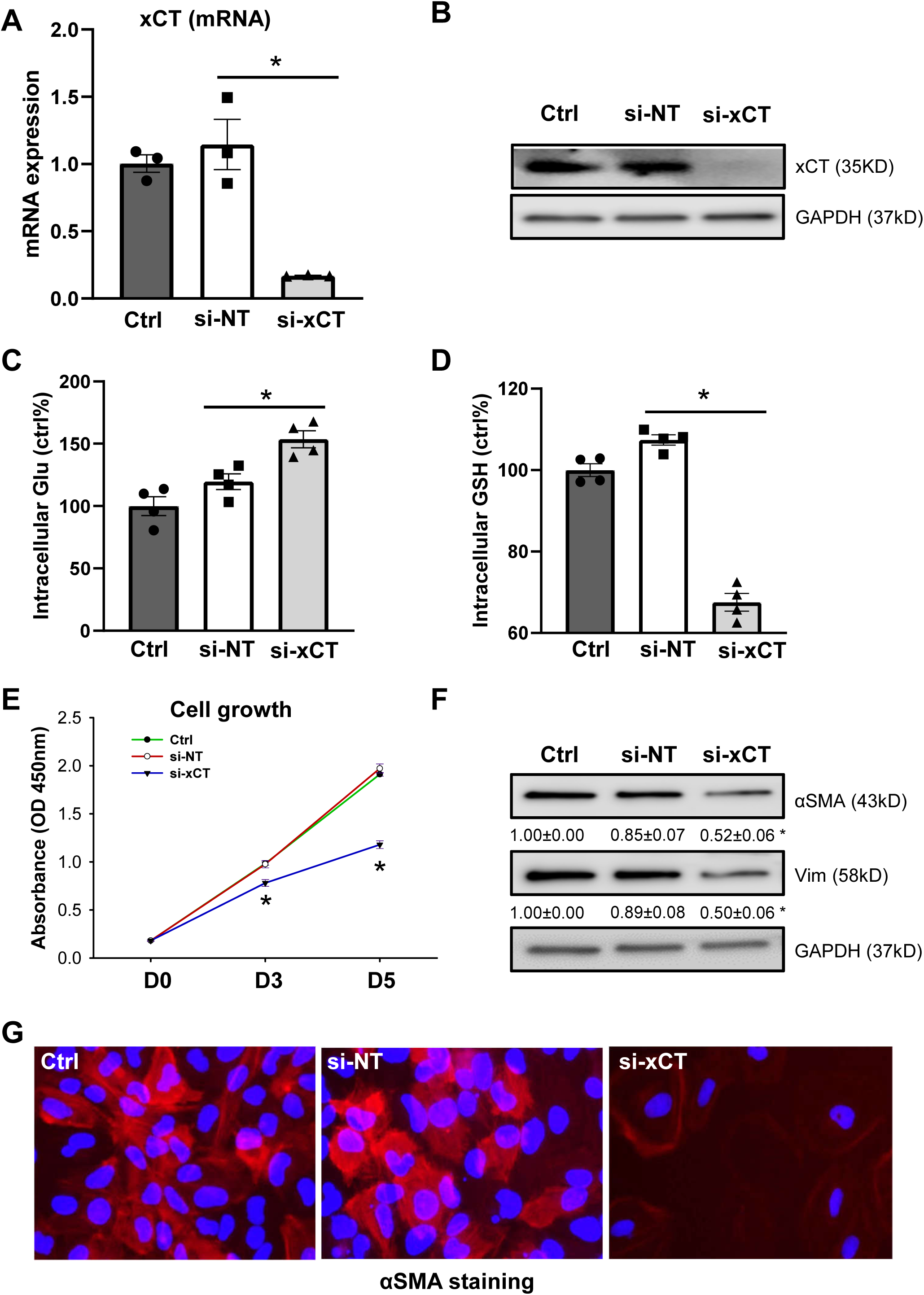
xCT deficiency suppressed growth and fibrogenic activity of MF-HSCs. Human MF-HSCs (LX2 cells) cells were treated with 20nM non-targeting siRNA (si-NT) or xCT siRNA (si-xCT). At day 5, **(A)** xCT mRNA was quantified by qRT-PCR; **(B)** xCT protein was quantified by western blotting; **(C)** Intracellular glutamate and **(D)** GSH were quantified and expressed as % of control level; **(E)** Cell growth was determined by CCK8 assay; **(F)** Protein expression of αSMA and vimentin was quantified by western blotting with GAPDH as the loading control. **(G)** αSMA expression was determined by immunocytochemistry (ICC). Bars represent mean ± SEM of n = 3-5 assays. *p < 0.05 vs non-targeting RNA group.

Pharmaceutical inhibitors of xCT, erastin or sulfasalazine (SASP),^5^ were also used to evaluate the role of xCT in human MF-HSCs. Consistent with the findings using siRNA, both xCT inhibitors suppressed MF-HSC growth in a dose-dependent manner (**Fig.2A; Suppl. Fig. 2A)**. xCT inhibition also increased intracellular glutamate (**Fig.2B**) and reduced GSH (**Fig.2C**), indicating the functional efficacy of erastin. We observed that inhibiting xCT also decreased expression of the MF-HSC differentiation markers αSMA and VIM and decreased expression of Col1α1 and the extracellular matrix remodeling factor MMP2, but increased the expression of quiescent marker PPARγ **(Figs. 2D, E).** In another MF-HSC cell line (rat 8B cells), inhibiting xCT by either erastin or SASP also dose-dependently inhibited MF-HSC growth **(Suppl. Fig. 2B, C)** and expression of αSMA, Col1α1 and MMP2 with an increase in the quiescence marker PPARγ **(Suppl. Fig. 2D, E).** Together, these data strongly indicate that xCT is critically required for MF-HSC to grow and maintain myofibroblastic activity.

**Figure 2.**
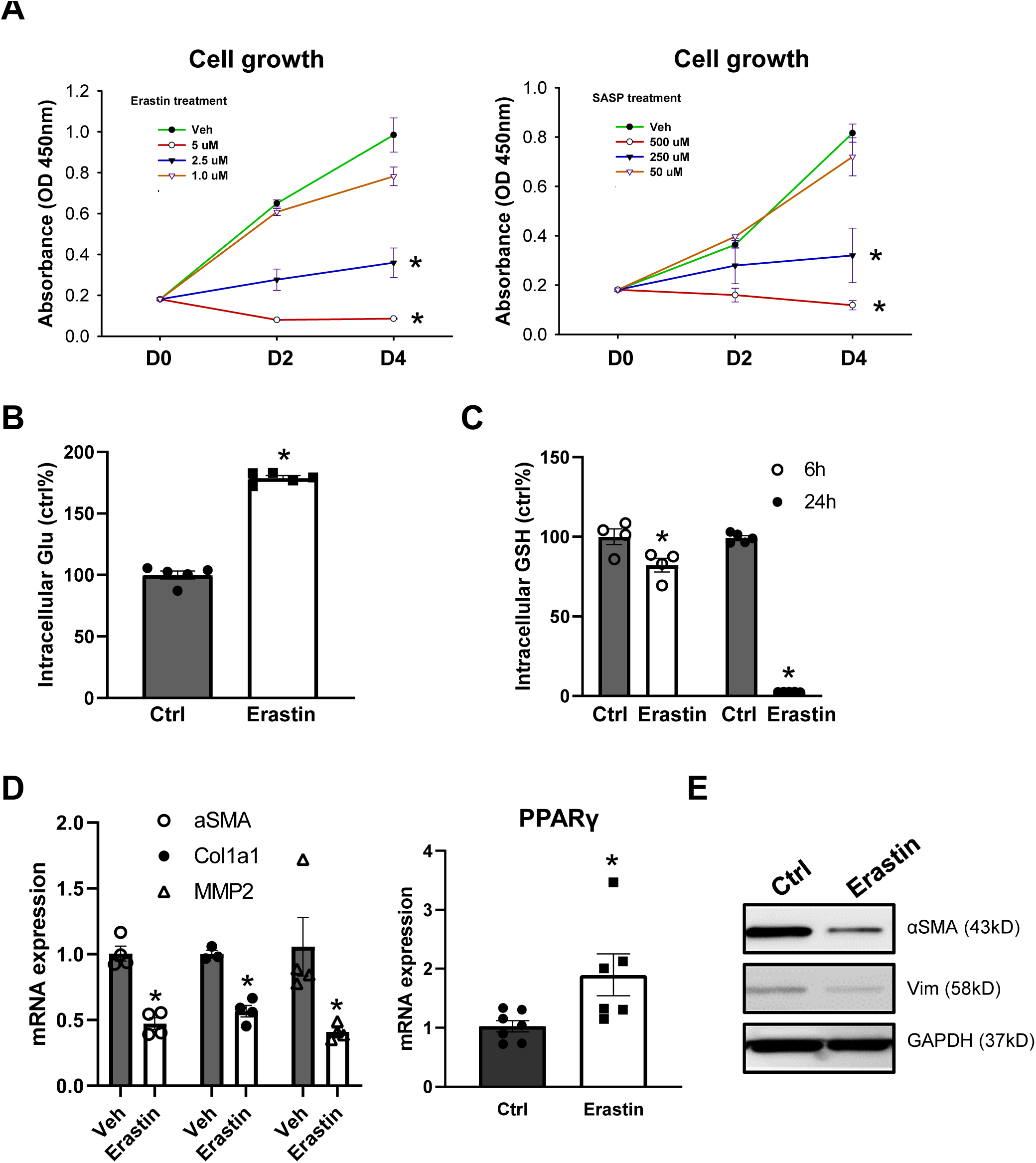
Pharmacologically inhibiting xCT suppressed growth and fibrogenic activity of MF-HSCs. Human MF-HSCs (LX2 cells) cells were treated with xCT inhibitors (erastin; SASP) or vehicle conrol (0.1% DMSO) for up to 4 days. **(A)** Cell growth was determined by CCK8 assay. **(B)** Intracellular glutamate level was quantified at 2h after 2.5 μM erastin treatment. **(C)** Intracellular GSH level was quantified at 6h and 24h after 2.5 μM erastin treatment. **(D)** mRNA was quantified by qRT-PCR. **(E)** Protein expression of αSMA and vimentin was quantified by western blotting with GAPDH as the loading control. Bars represent mean ± SEM of n = 3-5 assays. *p < 0.05 vs ctr group.

### Antioxidants rescue suppressed MF-HSC myofibroblastic activity during xCT inhibition

Cystine imported by xCT is immediately reduced to cysteine for biosynthesis of GSH, which is widely known as the first-line anti-oxidative defense. Thus, we hypothesized that the suppressed myofibroblastic activity induced by xCT inhibition is at least partially mediated by reduced GSH and/or increased oxidative stress. To test this hypothesis, we supplemented the erastin- or SASP-treated human MF-HSC cells with either the GSH precursor (N-acetylcysteine, NAC), or β-mercaptoethanol (β-ME, which can reduce extracellular cystine to cysteine and thus bypass the requirement of xCT since cysteine can be taken up by other transporters) to replenish GSH stores.^5, 6^ Both NAC and β-ME almost fully rescued the reduced MF-HSC viability caused by SASP or erastin in either the LX2 cells **(Figs. 3A, B, C)** or 8B cells **(Suppl. Fig. 3A, B; Suppl. Fig. 4A, B).** The suppressed expression of MF-HSC markers, such as αSMA, Col1α1 and VIM, was also rescued by NAC and β-ME in human MF-HSC (**Figs. 3D, E**) and rat MF-HSC cells (**Suppl. Fig. 4C-G**). Collectively, these data indicate that the suppression of myofibroblastic activity caused by xCT inhibition is mediated by GSH depletion and resulting oxidative stress.

**Figure 3.**
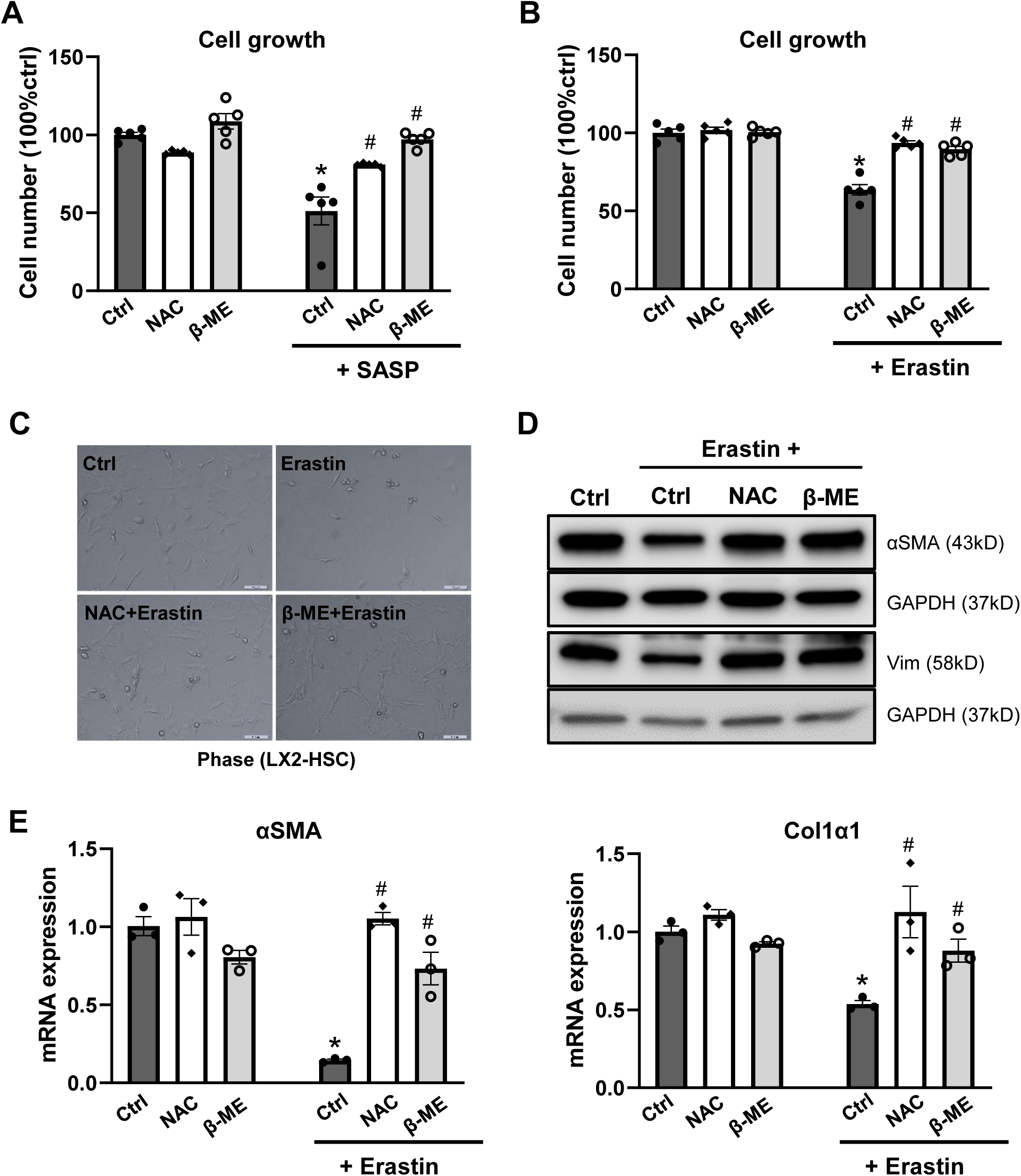
Suppressed myofibroblastic activity of MF-HSC caused by xCT inhibition was rescued by antioxidants. Human MF-HSCs (LX2 cells) cells were treated with xCT inhibitor erastin (2 μM), SASP (250 μM) or vehicle ctr (0.1% DMSO) for up to 4 days. In some experiments, the culture medium was supplemented with 2 mM NAC or 50 μM β-mercaptoethanol (β-ME). **(A, B)** Cell growth was determined at day 4 by CCK8 assay. **(C)** Representative phase pictures of LX2 cells. **(D)** Protein expression of αSMA and vimentin was quantified by western blotting with GAPDH as the loading control. **(E)** mRNA of αSMA and Col1α1 was quantified by qRT-PCR. Bars represent mean ± SEM of n = 3-5 assays. *p < 0.05 vs ctr group. ^#^p < 0.05 vs erastin or SASP alone group.

### xCT inhibition induces MF-HSC ferroptosis

xCT mediated-cystine uptake supports GSH biosynthesis and protects cells from oxidative stress and ferroptotic cell death.^13^ Therefore, we hypothesized that xCT inhibition induces ferroptosis in MF-HSCs. In support of this hypothesis, we found that erastin caused dramatic cell death, as indicated by the direct measurement of the DNA debris released into the culture medium (**Fig. 4A);** increased lipid peroxidation (C11 BODIPY assay) (**Fig. 4B**); and induced the expression of ferroptosis markers (*CHAC1* and *PTGS2*)^15, 16^ (**Fig. 4C**). Consistent with these findings, knocking down xCT expression by siRNA also induced the mRNA expression of ferroptosis markers (*CHAC1* and *PTGS2*) **(Suppl. Fig. 5A)**. Importantly, the canonical ferroptosis inhibitor ferrostatin and the iron chelator deferoxamine (DFO) almost fully reversed the increased lipid peroxidation (**Suppl. Fig. 5B**), rescued the reduced cell viability **(Figs. 4D, E, F; Suppl. Fig. 5C),** and restored expression of myofibroblastic markers **(Figs. 4G, H, I)**. In contrast, neither the apoptosis inhibitor Z-VAD-FMK, nor the necroptosis inhibitor Necrostain-1, improved cell viability **(Figs. 4F; Suppl. Fig. 5C)**.

**Figure 4.**
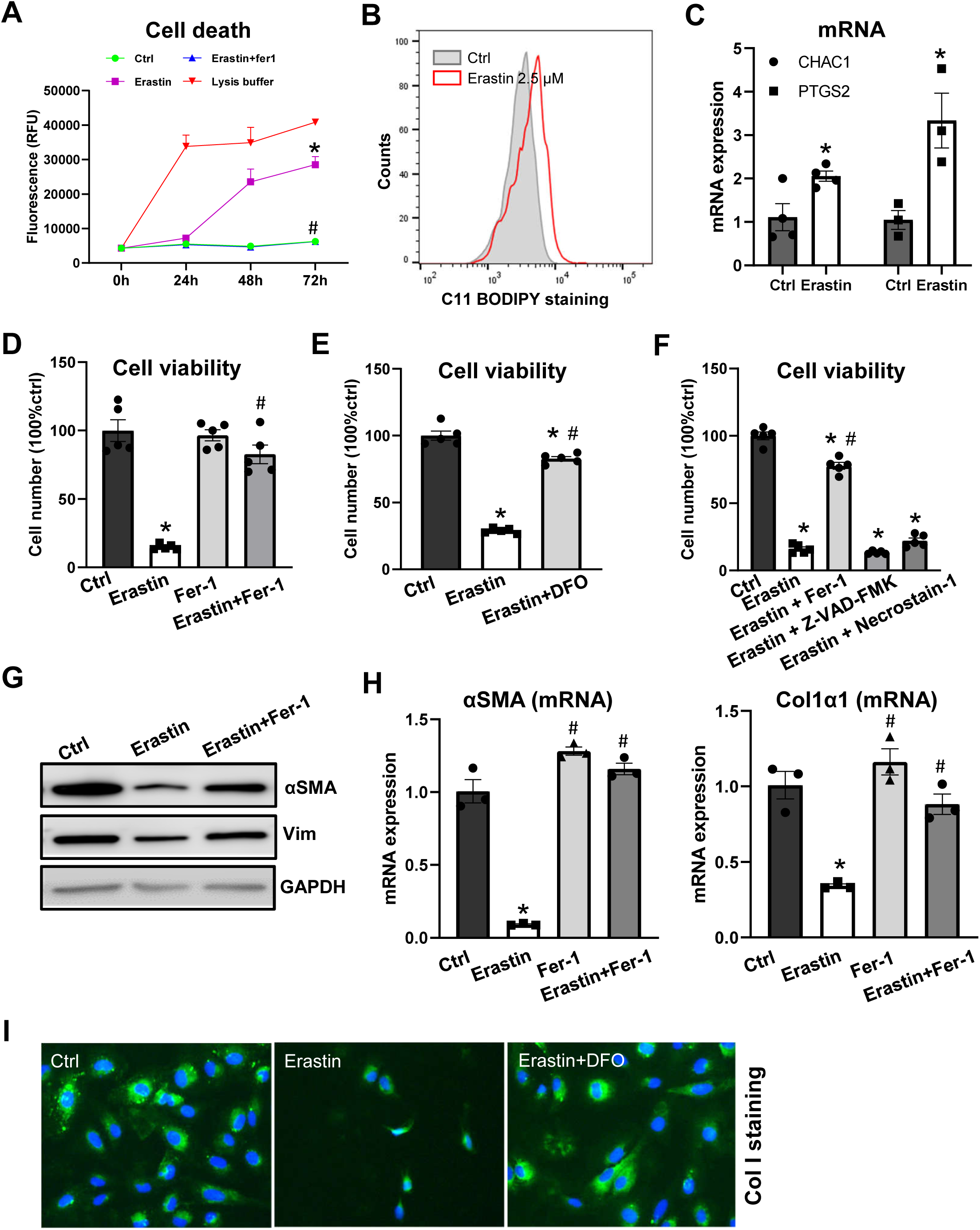
xCT inhibition induced MF-HSC ferroptosis. Human MF-HSCs (LX2 cells) cells were treated with xCT inhibitor erastin (2.5 μM) or vehicle ctr (0.1% DMSO) for up to 3 days. In some experiments, the culture medium was supplemented with the canonical ferroptosis inhibitor ferrostatin-1 (Fer-1) (2 μM), iron chelator deferoxamine (DFO) (10 μM), apoptosis inhibitor Z-VAD-FMK (10 μM) or necroptosis inhibitor Necrostain-1 (0.5 μM). **(A)** Cell death was assessed using CellTox™ Green Cytotoxicity assay. Cell lysis buffer was included as the positive control of complete cell death. **(B)** Lipid ROS was assessed by C11-BODIPY assay. **(C)** mRNA of CHAC1 and PTGS2 was quantified by qRT-PCR at day 3. **(D, E, F)** Cell growth was determined by CCK8 assay. **(G)** Protein expression of αSMA and vimentin was quantified by western blotting with GAPDH as the loading control. **(H)** mRNA of αSMA and Col1 was quantified by qRT-PCR. **(I)** Protein expression of Col1 was determined by ICC. Bars represent mean ± SEM of n = 3-5 assays. *p < 0.05 vs ctrl group. ^#^p < 0.05 vs erastin group.

Furthermore, erastin did not induce caspse-3 activation, a standard marker for apoptosis **(Suppl. Fig. 5D)**, while agents that replenish GSH stores (NAC and β-ME) blocked increased expression of the ferroptosis marker *CHAC1* **(Suppl. Fig. 5E)**. To examine whether the suppressed myofibroblastic phenotype is a direct effect of xCT inhibition or an indirect effect of increased cell death, we harvested viable HSCs 18h after erastin treatment, when cell death is still minimal **(Suppl. Fig. 6A)**. Interestingly, we found that short-term xCT inhibition is sufficient to cause MF-HSC to revert to a less myofibroblastic state, as indicated by a dramatic decrease in expression of fibrogenic genes and the extracellular matrix remodeling factor MMP2 **(Suppl. Fig. 6B-D)**. Cell death dramatically increased when MF-HSCs were treated with xCT inhibitor for longer periods of time (e.g., > 24h) (**Fig. 4A**). Nevertheless, the cells that survive long-term xCT inhibition were also more quiescent and less fibrogenic (**Fig. 4G-I)**. Together, these data demonstrate that although prolonged xCT inhibition generally induces ferroptosis in MF-HSCs, short-term xCT inhibition exerts anti-myofibroblastic effects.

Since MAPK pathway activity is important in erastin-induced ferroptosis in cancer cells,^6^ we examined whether this is also involved in MF-HSC ferroptosis. Both SP600125 (an inhibitor of JNK phosphorylation) and SB202190 (an inhibitor of p38 activation) rescued the cytotoxicity induced by erastin in human LX2 cells **(Suppl. Fig. 7A, B, C)** and rat 8B cells **(Suppl. Fig. 7D, E)**, indicating that MAPK pathway is also critical for erastin-induced ferroptosis in MF-HSCs.

### Glutaminolysis derived glutamate is critical for xCT-mediated cystine uptake in MF-HSCs

Our previous study demonstrated that glutaminolysis-derived glutamate is converted into α-KG to replenish the TCA cycle for energy production and anabolism.^4^ To test whether some of the glutaminolysis-derived glutamate is also used for exchange of extracellular cystine via xCT, we measured intracellular GSH levels in cells treated with the GLS1-specific inhibitors, CB839 or BPTES.^17^ We found that inhibiting glutaminolysis significantly decreased intracellular GSH levels (**Suppl. Fig. 8A**); confirmed that glutaminolysis inhibition suppressed HSC growth and expression of the fibrogenic gene, aSMA; and showed that these noxious effects were partially rescued by the GSH precursor NAC (**Suppl. Fig. 8B, C, D, E, F**). Together, these results suggest that MF-HSC normally use some of their glutaminolysis-derived glutamate to drive the cystine import to fuel GSH production and limit oxidative stress. Interestingly, however, βME (which acts independently of glutamate by converting extracellular cysteine to cystine for xCT uptake) did not rescue either cell growth or aSMA expression when glutaminolysis was blocked (**Suppl. Fig. 8B-F**), confirming that glutaminolysis is a critical source of glutamate for myofibroblast growth^4^. Paradoxically, the noxious consequences of inhibiting glutaminolysis were exacerbated by adding erastin to BTPES, suggesting that the glutaminolysis-derived glutamate is also needed to drive cystine import via xCT for GSH biosynthesis and antioxidant defense. xCT inhibition induced-retention of intracellular glutamate confounds, rather than alleviates, the negative effects of glutaminolysis inhibition. Therefore, the glutaminolysis-derived glutmate promotes fibrogenesis in MF-HSCs by at least two mechanisms (**Suppl. Fig. 7G**, **Suppl. Fig. 8H, J, Suppl. Fig. 8I, K**). Further, simply treating MF-HSC with either siRNA or pharmaceutical inhibitors of xCT suppresses Gls1 expression (**Suppl. Fig. 9A, B**), while inhibiting glutaminolysis by either siRNA or pharmaceutical inhibitors of Gls1 increases xCT expression (**Suppl. Fig. 9C, D)**, suggesting that xCT and GLS-1 normally interact to coordinate glutamine utilization with anti-oxidant defense. These new findings complement and extend our previous results,^4^ leading us to conclude that inhibiting glutaminolysis by either siRNA or pharmaceutical inhibitors of Gls1 increases xCT expression (**Suppl. Fig. 9C**, DHSCs use glutaminolysis-derived glutamate to fuel the TCA cycle and meet the high bioenergetic and biosynthetic demands of their myofibroblastic phenotype, as well as to enable xCT to import extracellular cystine for improved GSH biosynthesis and antioxidant defense during this high-growth state. This interpretation is consistent with knowledge in the cancer biology field which indicates that (like MF-HSCs) some malignant cells are typically “glutamine addicted” and require glutamine to fuel multiple carcinogenic pathways, including those that critically mediate biosynthetic and antioxidant functions.^18–20^ Thus, cells with high glutaminolytic activity are generally ‘growth-advantaged’ because glutamine catabolism generates sufficient glutamate to fulfill increased cellular biosynthetic-bioenergetic requirements, and to match the demand for enhanced antioxidant defense.

### Primary HSCs and hepatocytes are differentially sensitive to ferroptosis induced by xCT inhibition

Next, we tested whether primary HSCs are also sensitive to xCT inhibition. Primary HSCs were isolated from healthy mice; a portion of the pooled isolate was processed immediately to obtain quiescent (Q) HSCs (D0) and the remaining cells were cultured for 7 days (D7) to induce trans-differentiation into MF-HSCs **(Suppl. Fig. 10A)**. RNA sequencing analysis of D7 versus D0 HSCs revealed that xCT expression increased dramatically when Q-HSCs trans-differentiate into MF-HSCs **(Suppl. Fig. 10A)**, and this was confirmed by qRT-PCR and western blotting (**Fig. 5A**). As we had noted in MF-HSC cell lines (8B cells and LX2 cells), treating primary mouse HSCs with erastin (5 μM) decreased viability (**Fig. 5B**), and increased the mRNA expression of the ferroptosis markers *Chac1* and *Ptgs2* (**Fig. 5C),** suggesting that xCT inhibition also caused ferroptosis in primary HSC. Importantly, primary HSCs that survived erastin were less myofibroblastic, as indicated by a more quiescent morphology (**Fig. 5E**) and lower mRNA and protein expression of the MF-HSC markers, αSMA and Col1α1. (**Fig. 5D, E)**. To test whether quiescent HSCs are also sensitive to erastin, we treated the primary mouse HSCs with erastin or vehicle immediately after isolation, and harvested cells 3 days later. We found that xCT expression is rapidly induced in vehicle-treated HSCs even during early activation **(Suppl. Fig. 10B).** Hence, primary HSCs become sensitive to erastin treatment very early during the trans-differentiation process, as indicated by compromised net growth **(Suppl. Fig. 10C)**, higher rates of cell death **(Suppl. Fig. 10D)**, and selective survival of HSC with a more quiescent morphology **(Suppl. Fig. 10E)** and more quiescent gene expression profile **(Suppl. Fig. 10H)**. The aggregate data indicate that xCT is necessary for HSC to become myofibroblastic and to survive ferroptosis while maintaining the myogenic program characteristic of MF-HSCs.

**Figure 5.**
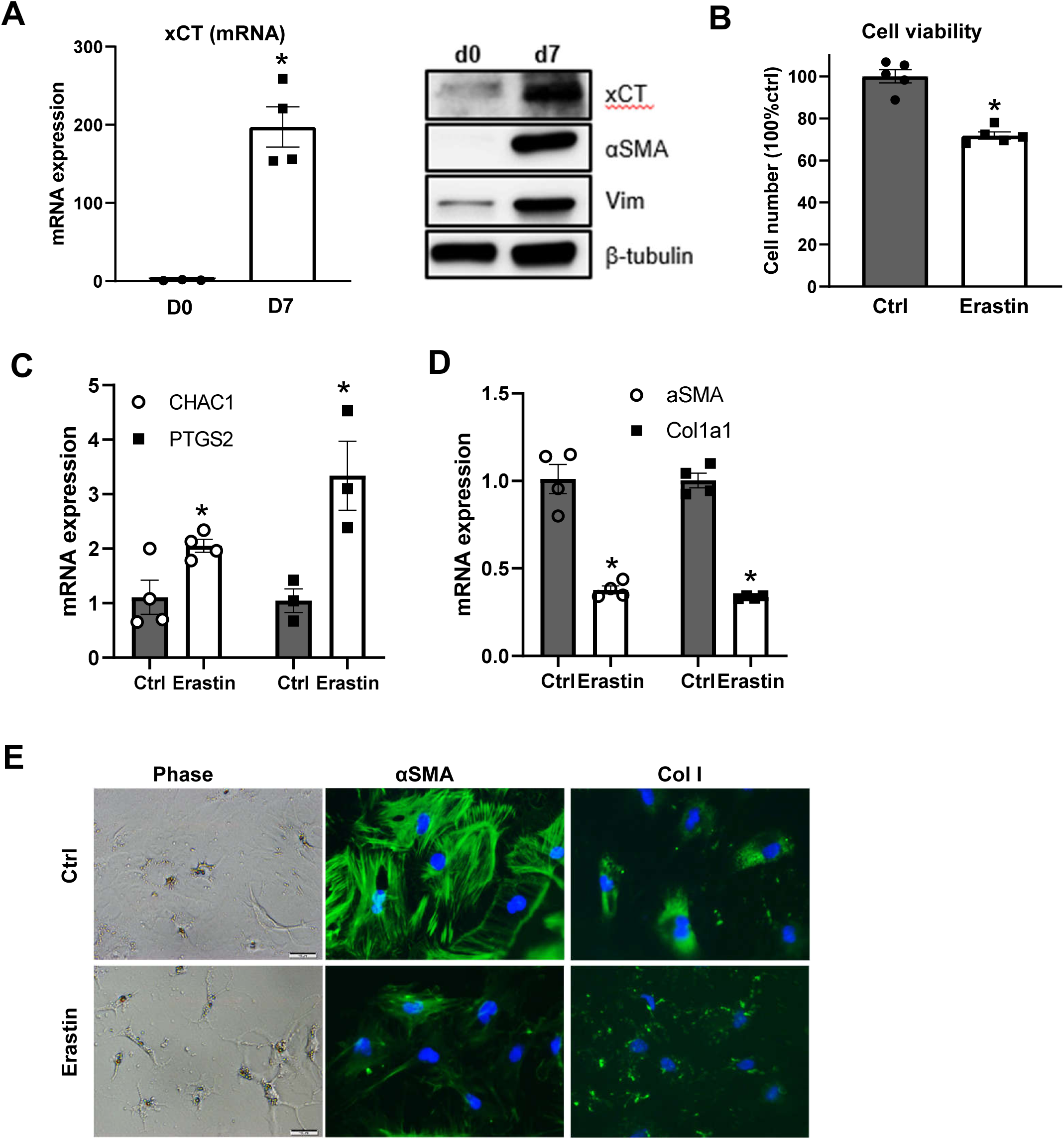
Primary HSCs are sensitive to xCT inhibition. **(A)** Primary HSCs were isolated from healthy C57Bl/6 male mice (n=4 mice/experiment). A portion of the pooled isolate was harvested for Q-HSCs (Day 0; D0) and remaining cells were cultured for 7 days to induce trans-differentiation. xCT expression was compared at D7 versue D0 by qRT-PCR and western blotting. **(B)** At day 4, some primary mouse HSCs were treated with erastin (5 μM) or vehicle ctr (0.1% DMSO). On day 7, cell growth/viability was assessed by CCK8 assay. **(C, D)** mRNA of Chac1, Ptgs2, αSMA and Col1 was quantified by qRT-PCR. **(E)** Representative phase pictures and immunofluorescent staining of αSMA and Col1 at day 7. Bars represent mean ± SEM of n = 3-5 assays. *p < 0.05 vs D0 (panel A) or vehicle control group (panel B-D).

Glutathione peroxidase (GPX4) critically regulates ferroptosis by operating downstream of xCT to detoxify lipid hydroperoxides to lipid alcohols.^21^ To determine if MF-HSCs succumb to ferroptosis when GPX4 is inhibited, human MF-HSCs were treated with RSL3, a direct GPX inhibitor, for 3 days with or without the ferroptosis inhibitor ferrostatin or the iron chelator DFO. We found that RSL3 also increased cell death **(Suppl. Figs. 11A)**, strongly inhibited MF-HSC viability **(Suppl. Figs. 11B, C)**, induced expression of the ferroptosis marker *PTGS2* **(Suppl. Fig. 11D),** and suppressed expression of MF-HSC markers **(Suppl. Fig. 11D)**. All these effects were largely reversed by the ferroptosis inhibitor ferrostatin **(Suppl. Figs. 11A, B, C, D)**, confirming that MF-HSCs are generally susceptible to ferroptosis triggered by theinhibitors of both xCT and GPX4.

To evaluate how HSC susceptibility to ferroptosis compares to that of other liver cell types, we treated primary hepatocytes (pHep) with similar doses of erastin and RSL3 that induce ferroptosis in primary HSCs. Interestingly, the xCT inhibitor (erastin) does not decrease pHep survival or increase expression of the ferroptosis markers *Chac1* and *Ptgs2* when used at a dose that causes robust ferroptosis in MF-HSCs, and the effects are still very minimal even at two-fold higher doses **(Suppl. Figs. 12A, B, C)**. In contrast, the GPX inhibitor (RSL3) dramatically decreases pHep viability and increases *Ptgs2* expression **(Suppl. Figs. 12A, B, C)**. In addition, the canonical ferroptosis inhibitor ferrostatin almost fully rescues compromised pHep viability **(Suppl. Figs. 12D)**. To determine the basis for these discordant findings, we compared expression of xCT and GPX4 mRNA in primary HSCs and pHep and discovered that expression of xCT transcripts is ∼80 fold higher in freshly isolated primary HSCs than in pHep **(Suppl. Figs. 12E)**. In contrast, GPX4 expression is robust in both cell types and higher in pHeps than HSCs **(Suppl. Figs. 12E)**. These results show that both HSCs and Heps are vulnerable to ferroptosis triggered by GPX4 inhibitor but indicate that cultured HSCs are more dependent upon xCT to maintain GSH stores and evade ferroptosis than cultured Heps. This is further supported by the finding that gene expression of critical enzymes involved in trans-sulfuration pathway is > 500 fold higher in primary mouse hepatocytes, and the rate-limiting enzymes in GSH biosynthetic pathway (gclc, gclm) is > 10 fold higher compared to primary HSCs **(Suppl. Figs. 12F)**. Together, these data demonstrate that hepatocytes mainly rely on the trans-sulfuration pathway for cysteine generation, and have a much more robust GSH biosynthetic capacity for antioxidant defense than HSCs. Because this differential sensitivity to xCT inhibition might simply reflect differences in the proliferative activity of the two cell types under their respective culture conditions, additional experiments were necessary to determine if HSCs and Heps respond differentially to xCT inhibition *in vivo*. The differential requirement for xCT in HSC and pHep may permit fibrogenic HSCs to be targeted without affecting pHep.

### xCT inhibition suppressed MF-HSC accumulation and liver fibrosis in acute liver injury

To determine whether xCT increases when HSCs are stimulated to transdifferentiate into MFs *in vivo,* an animal model of liver injury that rapidly induces both hepatocyte regeneration and MF-HSC accumulation was examined. Mice were injected with a single dose of CCl4, and then hepatocyte proliferative activity, liver injury, and MF-HSC accumulation were assessed at 48h after CCl4 treatment. We confirmed that CCl4 treatment acutely triggered death of peri-venular hepatocytes and induced proliferation of neighboring hepatocytes as well as massive accumulation of cells expressing the MF-HSC marker αSMA **(Suppl. Fig. 13A)** together with a dramatic increase in xCT expression in total liver tissue (**Fig. 6A**). To determine the cellular source for this xCT induction, we isolated primary HSCs from mice 48h after CCl4 treatment and examined xCT expression in these cells. Data from both the DNA gel (data not shown) and RT-PCR showed that xCT is highly induced in HSCs that were activated *in vivo* (**Fig. 6B**). Co-immunofluorescence staining of liver sections from CCl4-treated mice co-localized xCT with the MF-HSC marker aSMA, confirming that MF-HSCs were the major source of xCT in CCl4-injured livers (**Fig. 6C**). Next, we determined how inhibiting xCT with erastin influenced responses to CCl4 injection. Consistent with our findings in primary hepatocytes, treating mice with the xCT inhibitor erastin did not affect hepatocyte injury, as indicated by both serum ALT and AST levels **(Suppl. Fig. 13B)** and assessment of H&E-stained liver sections **(Suppl. Figs. 13C)**. Hepatocyte regeneration was also comparable in mice that were treated with vehicle and erastin after CCl4 exposure (**Suppl. Figs. 13D, E**). In contrast, we found that protein levels of MF-HSC markers αSMA and desmin were dramatically decreased in erastin-treated mice, as assessed by immunostaining (**Fig. 6D).** The reduction in expression of these proteins was paralleled by a decrease in desmin and Vim transcripts (**Fig. 6E**). mRNA levels of Col1α1, Col3α1 and Col6α1 were also reduced (**Fig. 6F),** and decreased Col1α1 expression was confirmed by western blotting **(Suppl. Figs. 13F)**. Together, these data suggest that MF-HSCs are particularly vulnerable to xCT inhibition and demonstrate that inhibiting xCT suppresses the accumulation of MF-HSC and inhibits liver fibrosis without exacerbating acute CCl4-induced liver injury.

**Figure 6.**
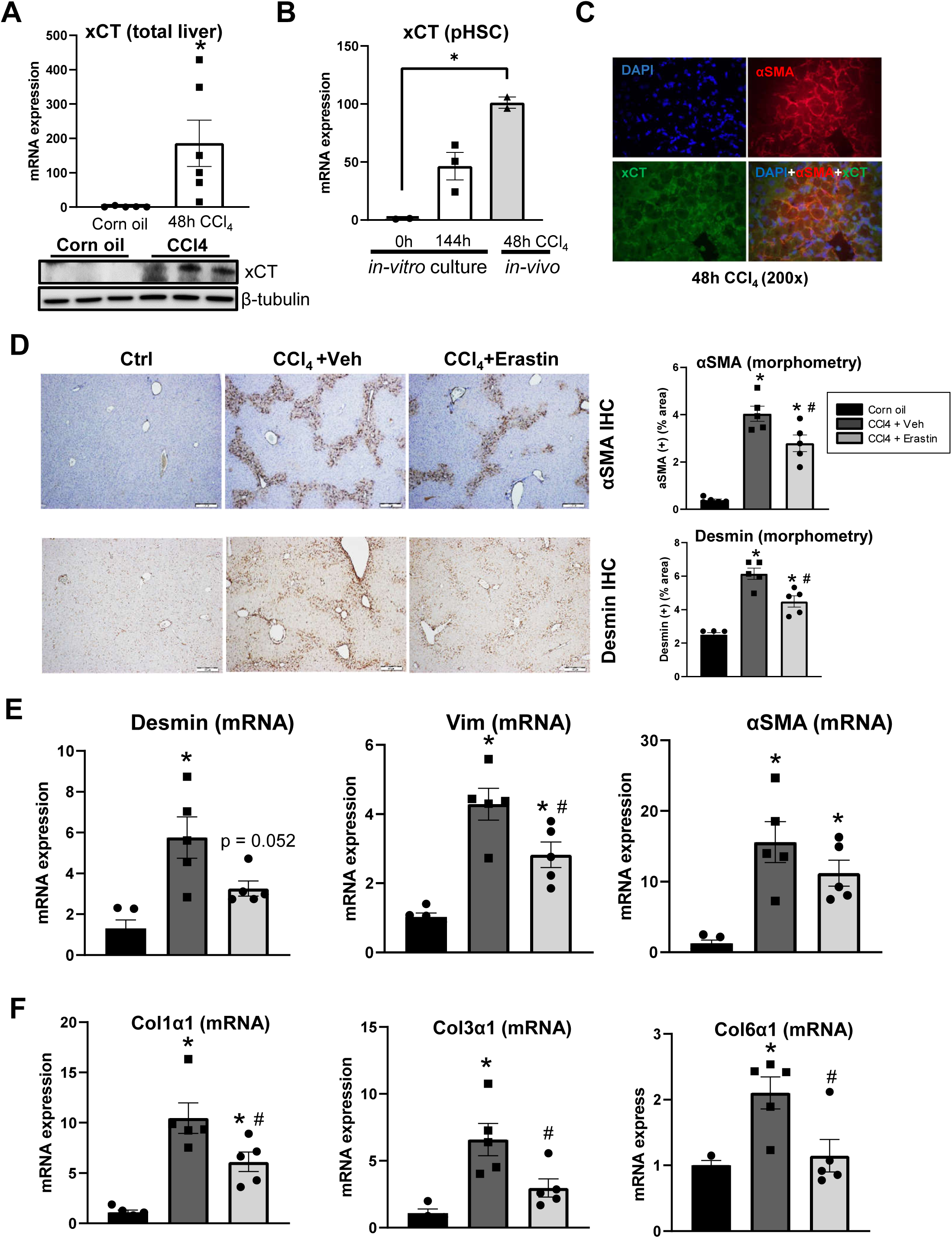
xCT inhibition suppressed MF-HSC accumulation and liver fibrosis in acute liver injury. Adult mice were injected intraperitoneally with corn oil vehicle or CCl_4_ (1200 mg/kg), liver tissues were harvested at 48h post-CCl_4_. **(A)** xCT expression was determined by qRT-PCR or western blotting. **(B)** Primary HSCs were isolated from healthy C57Bl/6 male mice or CCl_4_ treated mice and assayed immediately to compare expression of xCT. A portion of the cells from healthy mice were also cultured for 7 days to induce trans-differentiation. xCT expression was assessed by qRT-PCR. **(C)** Representative α-SMA and xCT co-immunofluorescence in liver of a CCl_4_-treated mouse. In subsequent experiments, some mice were intraperitoneally injected with erastin or its vehicle (10% DMSO) at 9h and 30h post-CCl4 injection, and liver tissues were harvested at 48h post-CCl4 (n=5 mice/group). **(D)** Representative α-SMA and desmin stained liver sections and corresponding densitometry analysis for stained (+) areas. **(E)** mRNA levels of MF-HSC marker α-SMA, desmin and vimentin determined by qRT-PCR. **(F)** mRNA levels of fibrogenic marker Col1α1, Col3α1 and Col6α1 determined by qRT-PCR. Bars represent mean ± SEM. *p < 0.05 vs corn oil. ^#^p < 0.05 vs CCl_4_+vehicle group.

## Discussion

Liver fibrosis develops when excessive hepatic stellate cells (HSCs) accumulate in the injured liver. While transient accumulation of MF-HSCs is necessary for effective liver regeneration, prolonged excessive accumulation of such cells leads to progressive fibrosis, defective repair, and ultimately, cirrhosis.^3^ Because the outcome of liver injury is dictated by factors that control the size of MF-HSC populations, the mechanisms that orchestrate MF-HSC accumulation are attractive therapeutic targets.^3^ However, therapies that block this process have yet to be discovered. In this study, we discovered that xCT is dramatically induced during HSC trans-differentiation and is critical for protecting MF-HSCs from ferroptosis. Consistent with this, xCT inhibition induced MF-HSC ferroptosis and suppressed liver fibrosis in the CCl4 acute liver injury model. In contrast, healthy hepatocytes express significantly lower levels of xCT and were relatively resistant to xCT inhibition following CCl4-induced acute hepatotoxicity. Further research is needed to determine if similarly beneficial results are obtained when xCT is inhibited in other types of liver injury because hepatocytes may up-regulate their expression of xCT to protect themselves from oxidative stress during chronic liver injury,^9^ narrowing the therapeutic window for this anti-fibrotic approach.

Strong experimental data indicate that oxidative stress plays a critical role in triggering liver damage and initiating liver fibrogenesis.^22^ However, the relationship between oxidative stress and HSCs is somewhat complex.^23^ While ROS and their products are known to stimulate activation and proliferation of HSCs, there is also compelling evidence that ROS induce HSC death.^23^ It was proposed that quiescent HSCs have a high GSH content that protects them from the detrimental effects of ROS, but that GSH levels decrease when HSCs transdifferentiate into MF-HSCs, rendering these cells vulnerable to ROS-induced death.^23^ Our experimental data provide important proof for such a concept. Therefore, elimination of activated HSCs through selective oxidative stress induction is plausible and might provide novel anti-fibrotic therapies for chronic liver disease. Cysteine is well-known to play pivotal roles in cellular redox homeostasis by serving as the rate-limiting amino acid for biosynthesis of GSH and redox-reactive proteins involved in anti-oxidant defense. By importing extracellular cystine to replenish intracellular cysteine pools, xCT has been shown to be essential for GSH synthesis and maintaining intracellular redox homeostasis.^13^ There is growing evidence that xCT expression is increased in various malignancies.^13^ HCC strongly expresses xCT and although xCT is barely detectable in healthy adult liver, it is induced by conditions that deplete glutathione and/or increase oxidative stress, ^7–9^ supporting the concept that xCT also has an important anti-oxidant defense role in nonmalignant liver. Oxidative liver injury stimulates HSCs to trans-differentiate into fibrogenic myofibroblasts and previously, we showed that growth of MF-HSC populations depends upon their increased glutaminolytic activity.^4^ Glutamine catabolism generates glutamate and xCT exports intracellular glutamate in order to import extracellular cystine for GSH synthesis, but whether or not xCT is induced during HSC trans-differentiation and how this affects MF-HSC accumulation and liver fibrosis were virtually unknown. Our current study provides novel evidence that xCT is dramatically induced during HSC trans-differentiation and shows that MF-HSCs require xCT to remain proliferative and myofibroblastic, linking their metabolic adaptations with enhanced resistance to ferroptosis. Indeed, inhibiting xCT in MF-HSCs by either siRNA or inhibitors blocked intracellular GSH biosynthesis and led to suppressed cell growth and fibrogenic gene expression. Conversely, supplementing culture medium with NAC or β-ME, factors that replenish GSH stores, rescues the inhibited myofibroblastic activity caused by xCT inhibition. More importantly, the xCT inhibitor erastin also decreased MF-HSC accumulation in the CCl4-induced acute liver fibrosis model without affecting hepatocyte injury. These finding clearly demonstrate that xCT is critically required for MF-HSC accumulation during liver injury and thus may represent an attractive therapeutic target to treat liver fibrosis.

Previous research indicates that mechanisms that restrict contractility, inhibit energy metabolism, and induce senescence in cultured MF-HSCs inhibit fibrogenesis and thus, are potentially attractive approaches to reduce fibrosis in chronic liver diseases.^3^ In addition, inducing HSC cell apoptosis to promote fibrosis resolution has been proposed as a treatment for cirrhosis based on evidence that increased HSC apoptosis accompanies resolution of fibrosis in pre-clinical models.^24, 25^ However, human HSCs are relatively resistant to apoptosis and it is not yet clear if increased apoptosis of MF-HSCs occurs when fibrosis regresses after liver injury resolves in humans.^24, 26, 27^ It is also uncertain whether apoptosis can be increased selectively in HSCs without reducing hepatocyte viability and worsening liver function in the context of chronic liver injury. Although caspase-independent (i.e., non-apopototic) pathways of HSC death have been described, very little work has been done to examine if HSC death could be increased selectively by targeting these other pathways for programmed cell death.^28–30^ Improved understanding of how non-apoptotic cell death pathways act in HSC may allow more specific and efficient anti-fibrotic therapies.

Recent studies suggest a complex role of ferroptosis in organ fibrosis which is influenced by cell type and context. Ferrostatin-1, the canonical ferroptosis inhibitor, was shown to suppress cardiac fibrosis by maintaining mitochondrial function.^31^ Liproxstatin-1, a more potent ferroptosis inhibitor, was shown to alleviate radiation-induced lung fibrosis by down-regulating TGF-β1.^32^ In murine models of oxidative stress caused by hepatocyte iron overload, ferrostatin-1 inhibited liver fibrosis by limiting hepatocyte injury.^33^ In contrast, our results suggest that globally inhibiting ferroptosis in other types of liver injury might increase liver fibrosis by enhancing accumulation of fibrogenic MF-HSCs. While our new findings may seem counter-intuitive, they are predicted by results of our studies in cultured liver cells, which clearly demonstrated that HSCs must inhibit ferroptosis in order to be myofibroblastic. It is important to point out that our results are consistent with many studies showing mesenchymal differentiation is associated with increased ferroptosis sensitivity ^34^ and cystine addiction. ^35^ Further, by directly manipulating the activity of various components of the ferroptotic pathway in cultured HSCs, we proved that MF-HSCs depend upon xCT to protect themselves from ferroptosis and maintain myofibrogenic activities. Consistent with those *in vitro* data, administering erastin to inhibit xCT activity in mice with acute toxic liver injury rapidly induced ferroptosis and immediately blocked accumulation of fibrogenic MF-HSCs. Interestingly, erastin did not exacerbate CCl4-induced acute liver injury, suggesting that hepatocytes might be less vulnerable to xCT inhibition than HSCs in that model. Indeed, we discovered that expression of xCT mRNA is almost a log-fold lower in primary hepatocytes than in primary HSCs. Further, we found that doses of erastin that massively induced ferroptosis in primary HSC cultures did not reduce the viability of primary hepatocyte cultures. The aggregate data suggest that MF-HSCs may be more dependent on xCT to maintain levels of GSH that GPX4 requires to detoxify lipid peroxides than hepatocytes. It is interesting to note that Hippo transducers (YAP/TAZ) and ataxia telangiectasia mutated (ATM), factors previously shown to be critical for HSC trans-differentiation^36, 37^, were also recently found to be critical for ferroptosis^38–40^, providing a potential mechanistic link between MF-HSC differentiation and ferroptosis vulnerability. Conversely, the relative resistance of hepatocytes to ferroptosis caused by xCT inhibition is supported by recent studies of hepatocyte toxicity caused by acetaminophen overdose, which revealed that hepatocytes predominately rely on the trans-sulfuration pathway to supply cystine to the redox system, and only induce xCT to import extracellular cystine when the trans-sulfuration pathway is impaired and unable to fulfill demands for GSH synthesis.^7^ More research is needed to determine if this differential sensitivity to xCT inhibitors or other ferroptosis inducers can be exploited therapeutically to safely inhibit fibrogenesis in other types of liver injury. The tenet merits further evaluation in light of recent reports showing that artemether and artesunate, which were recently identified as ferroptosis inducers in HSCs, suppressed liver fibrosis.^41, 42^ In addition, sorafenib alleviated bile duct ligation (BDL)-induced liver fibrosis in mice and ferroptosis was shown to be induced in HSCs from fibrotic patients with HCC receiving sorafenib therapy.^43^ Given growing evidence that vulnerability to ferroptosis is tissue-, cell type, and/or context-dependent, novel anti-fibrotic approaches may be developed by exploiting these differences.

In summary, by deploying an array of complementary tools (human and rat MF-HSC cells lines, primary mouse HSCs and a CCl4-induced acute liver fibrosis model), we demonstrated that targeting xCT and/or ferroptosis is a promising therapeutic strategy for limiting MF-HSC accumulation during liver fibrosis. We discovered that xCT is dramatically induced during HSC trans-differentiation for GSH biosynthesis and protection from oxidative stress. Its inhibition induced MF-HSC ferroptosis and suppressed liver fibrosis. In contrast, healthy hepatocytes express significantly lower levels of xCT and are relatively resistant to xCT inhibition during acute liver injury. However, because hepatocytes may need to up-regulate xCT to survive chronic liver injury, further research is needed to determine if targeting xCT can selectively induce ferroptosis in MF-HSCs and safely inhibit liver fibrosis during chronic liver injury.

## Author Contributions

K.D., J.A.C. and A.M.D. conceived of the experiments. K.D., S.H.O., T.S. and W.H.Y performed experiments. K.D., W.H.Y, J.A.C. and A.M.D. analyzed data. K.D., J.A.C. and A.M.D. wrote the manuscript. Everyone reviewed and approved the manuscript.

---

**Suppl. Fig. 1.**
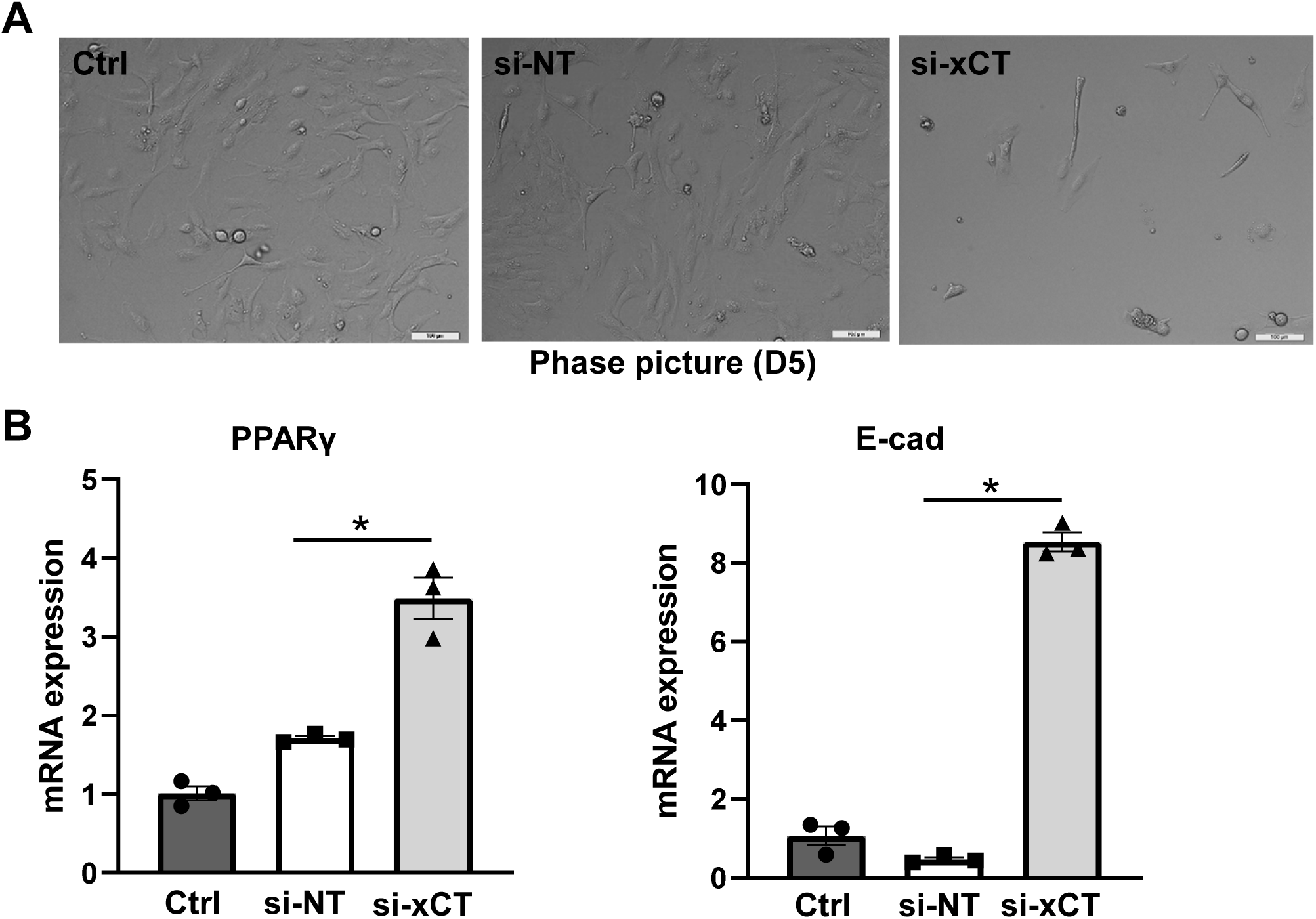

**Suppl. Fig. 2.**
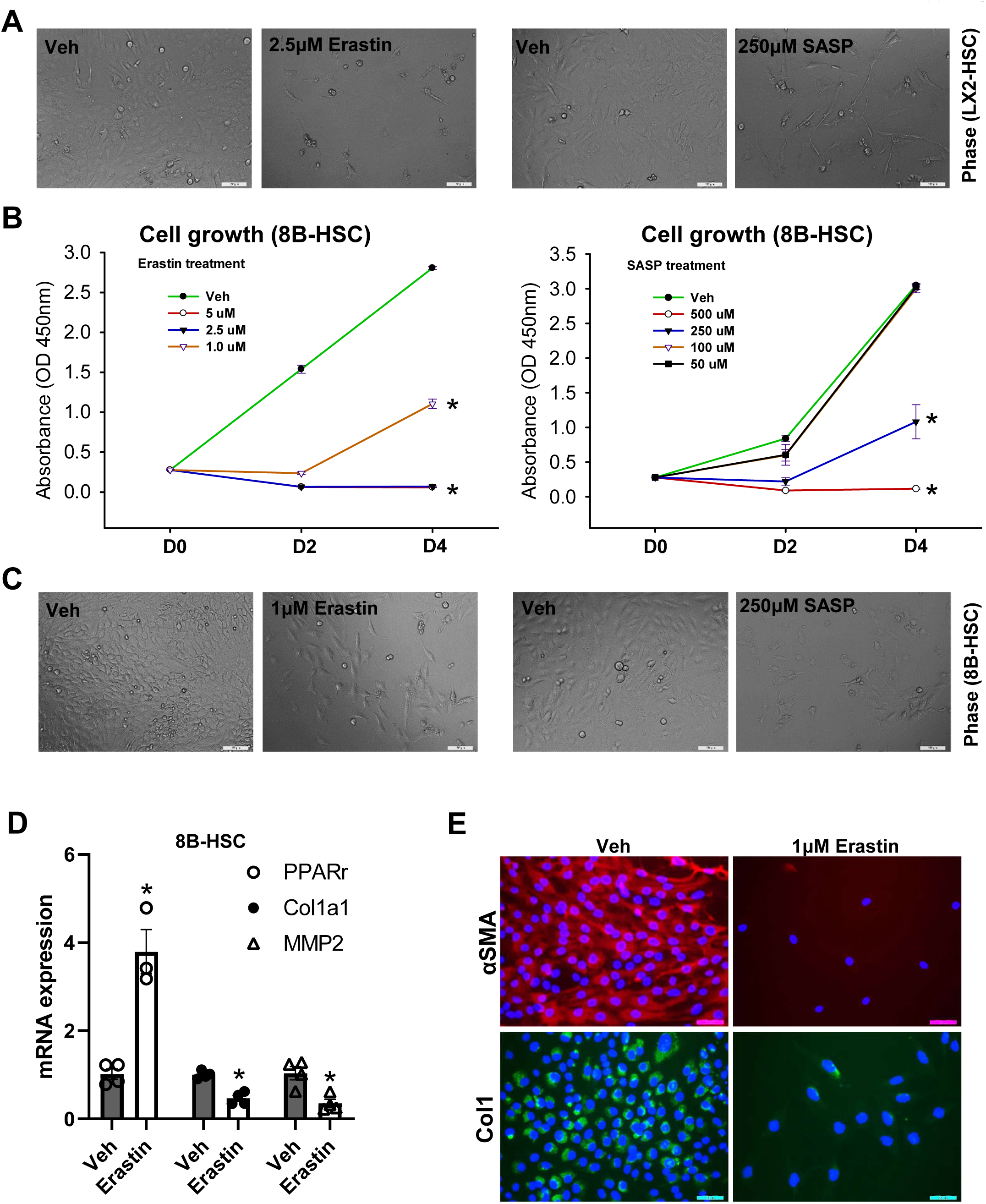

**Suppl. Fig. 3.**
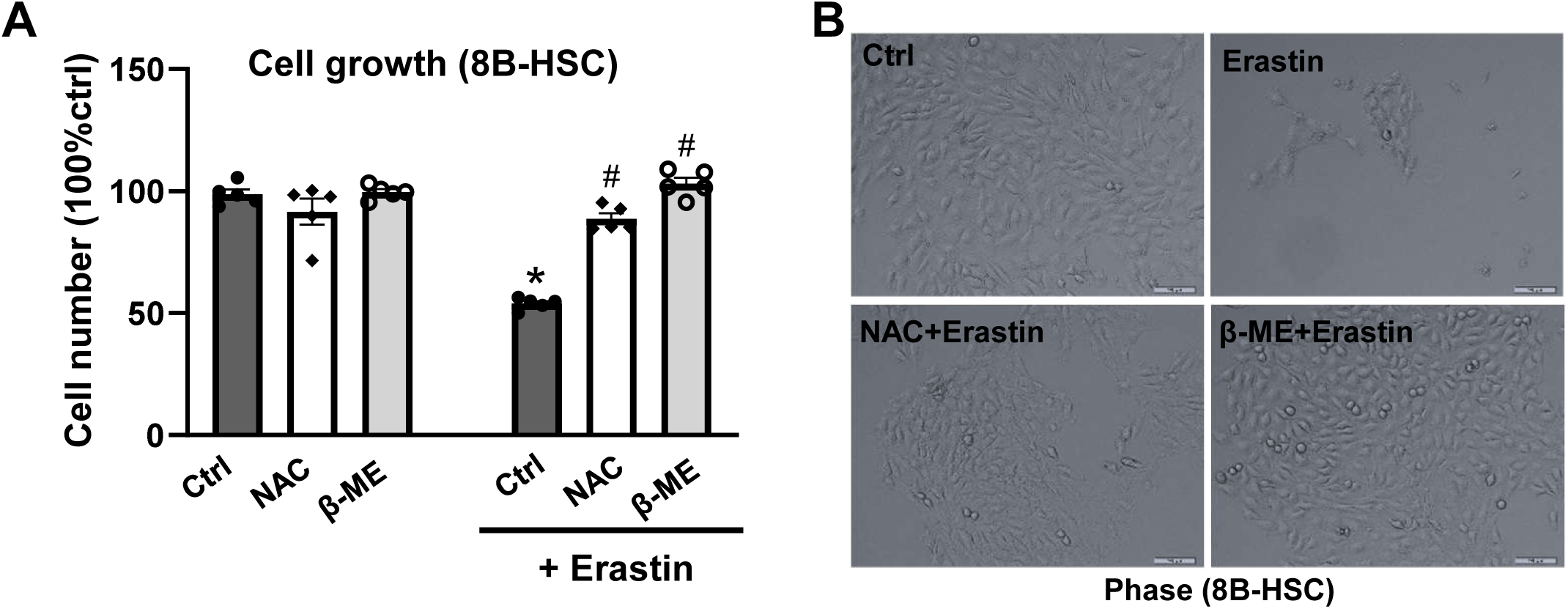

**Suppl. Fig. 4.**
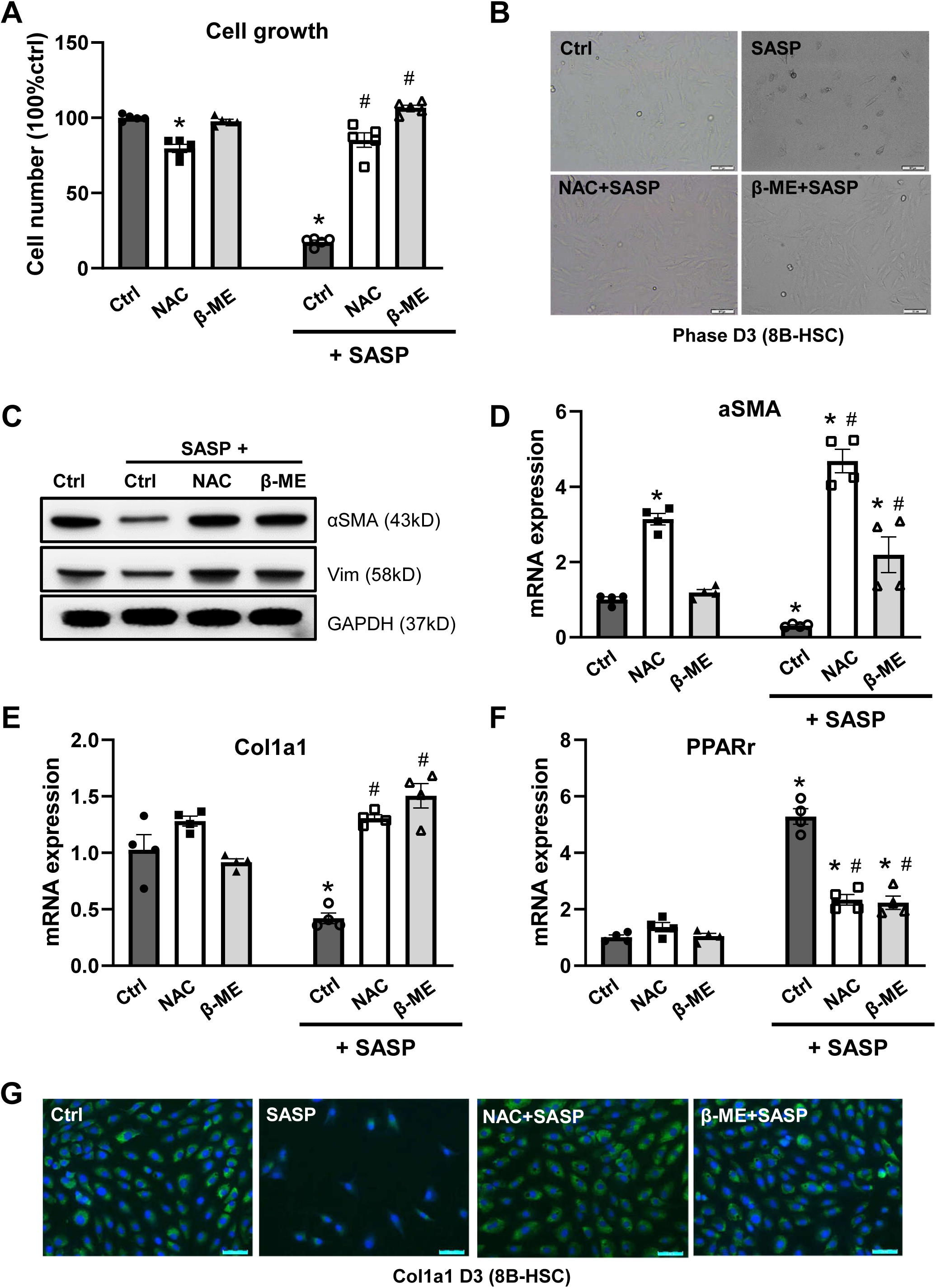

**Suppl. Fig. 5.**
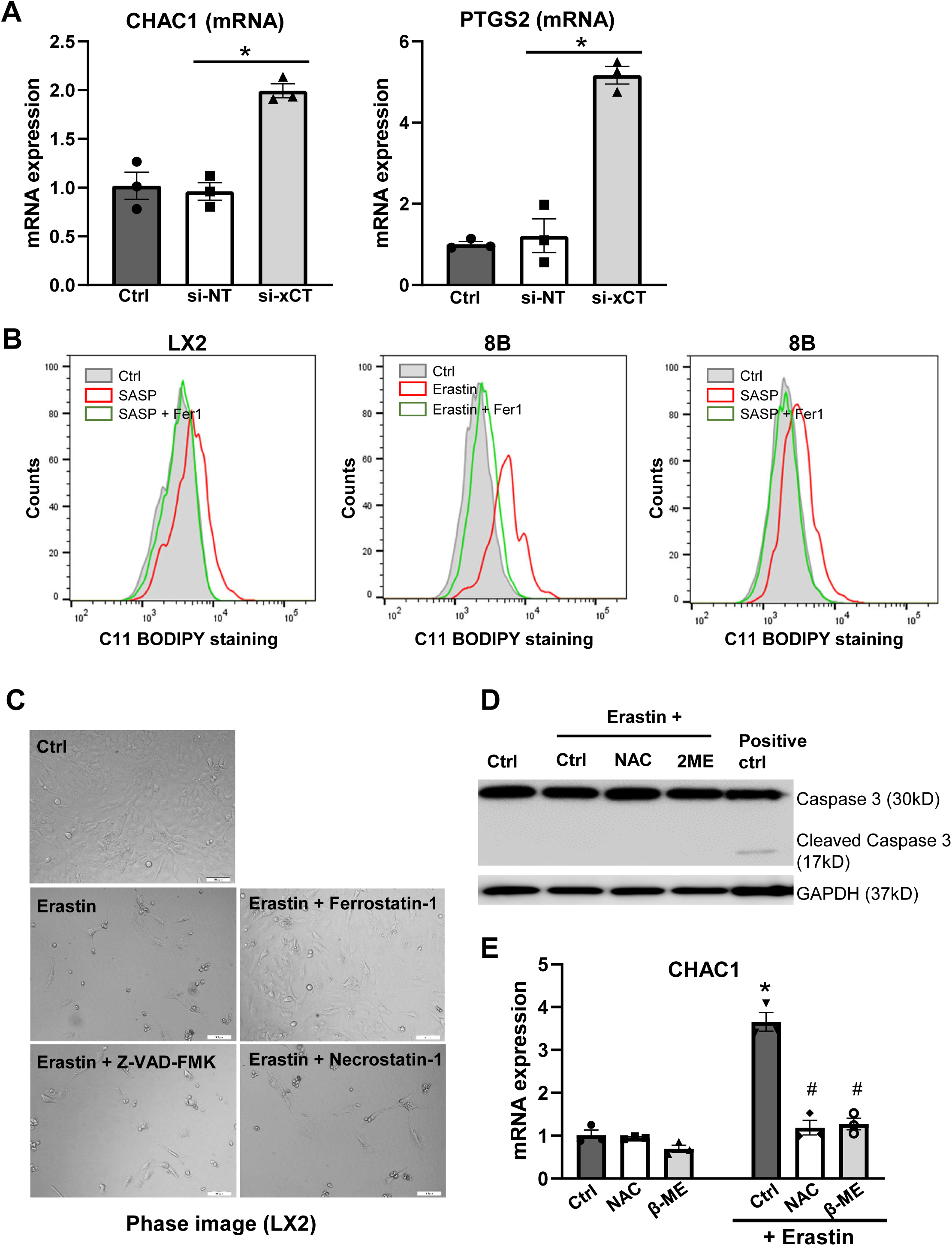

**Suppl. Fig. 6.**
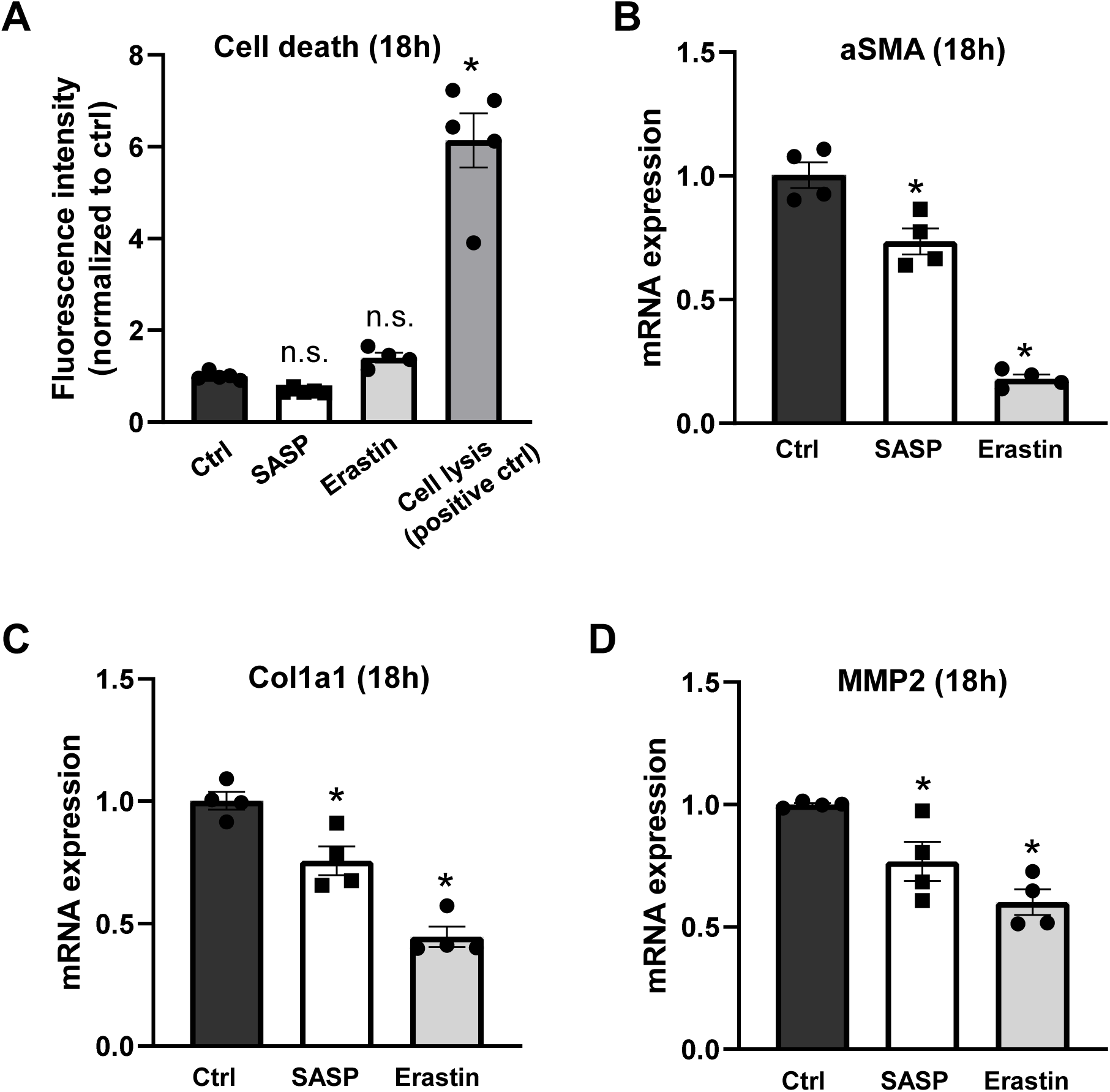

**Suppl. Fig. 7.**
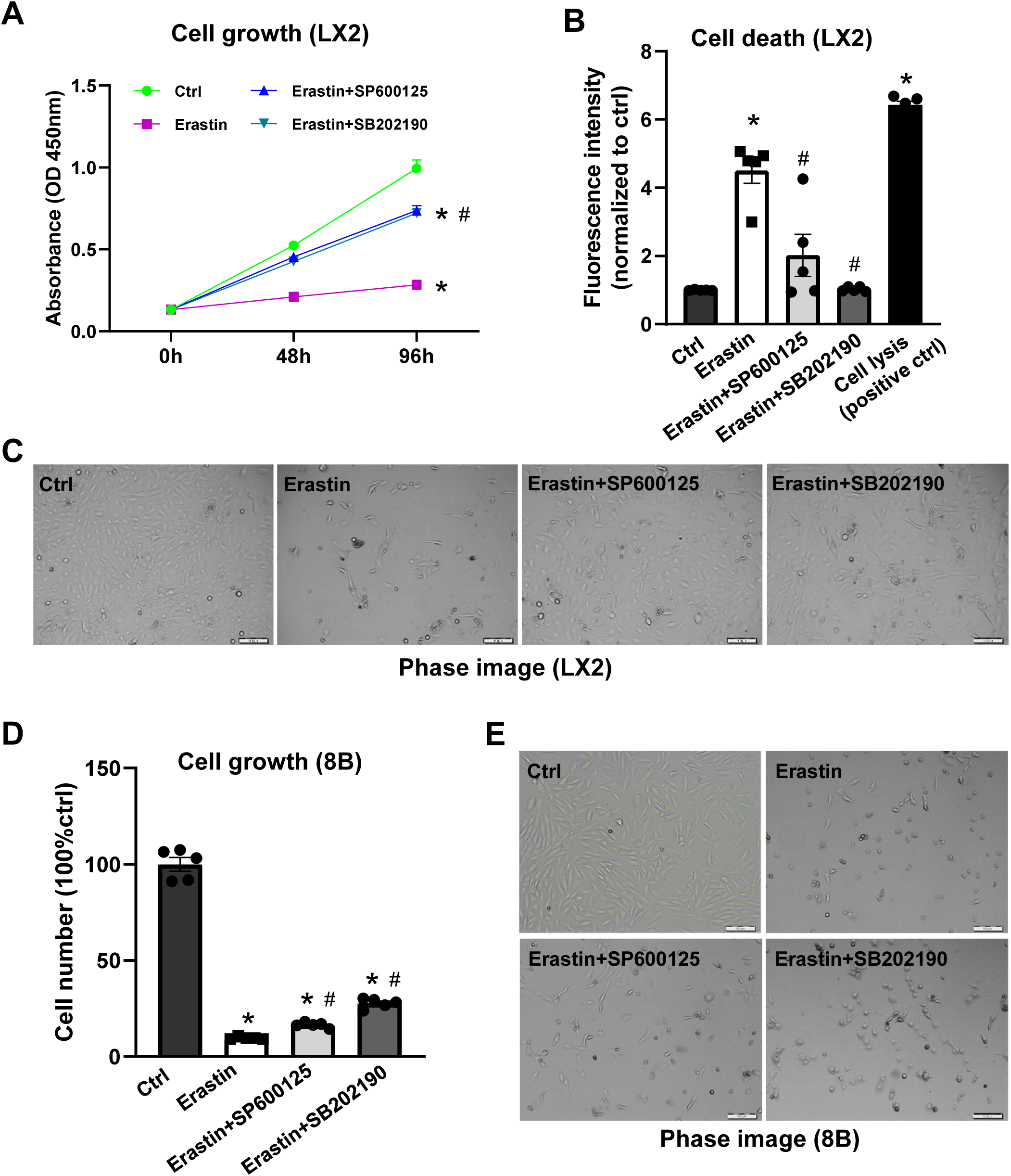

**Suppl. Fig. 8.**
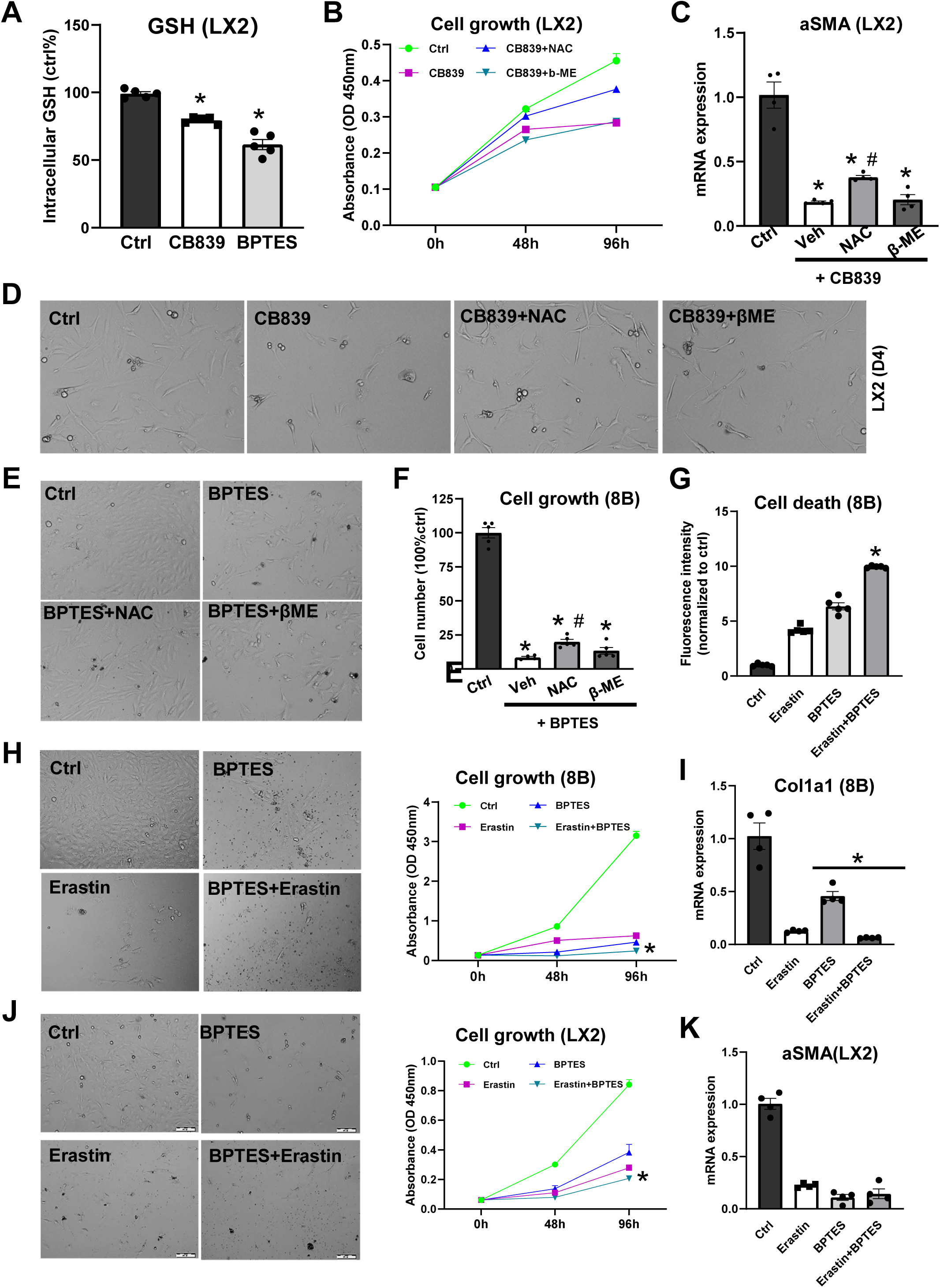

**Suppl. Fig. 9.**
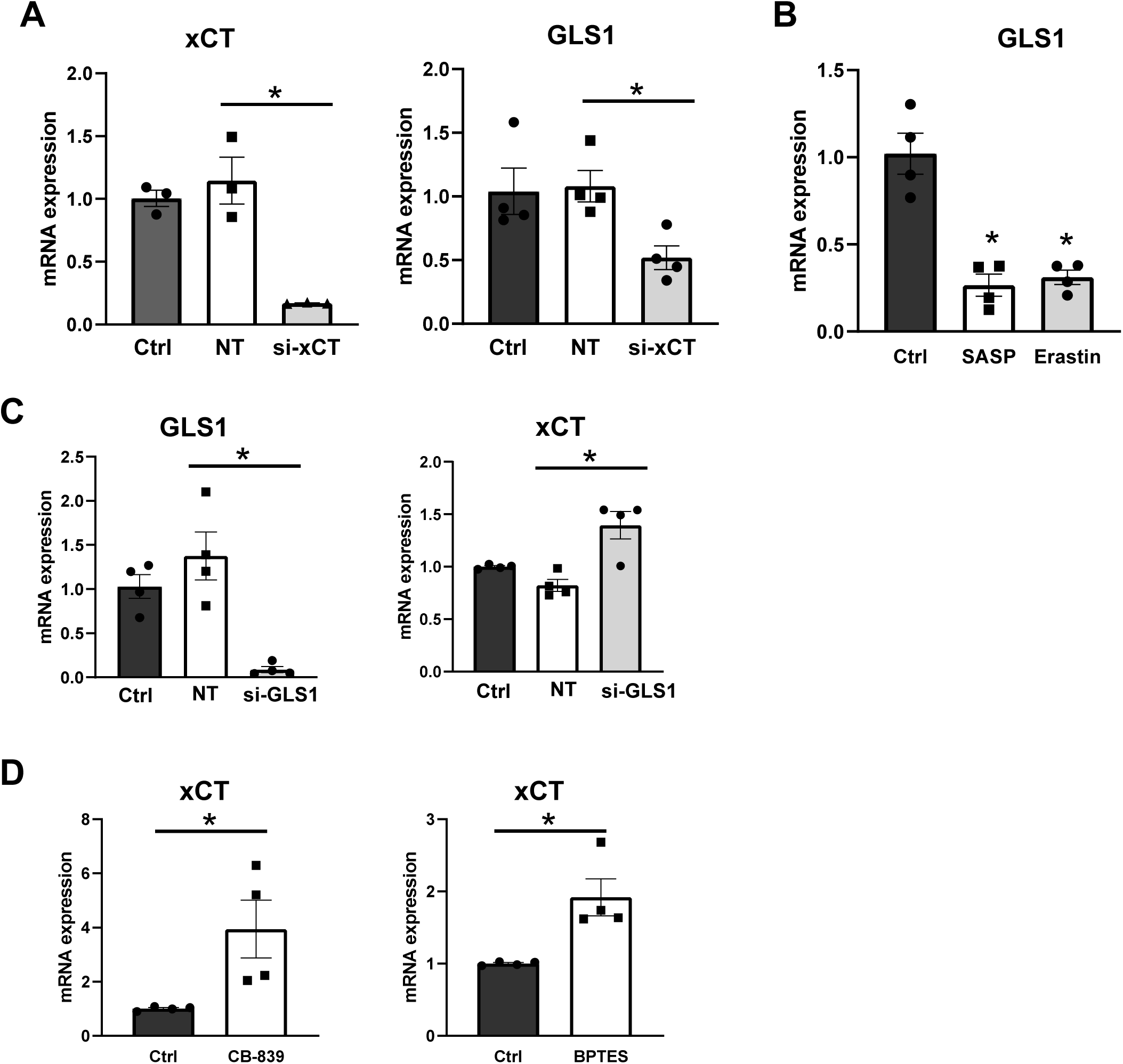

**Suppl. Fig. 10.**
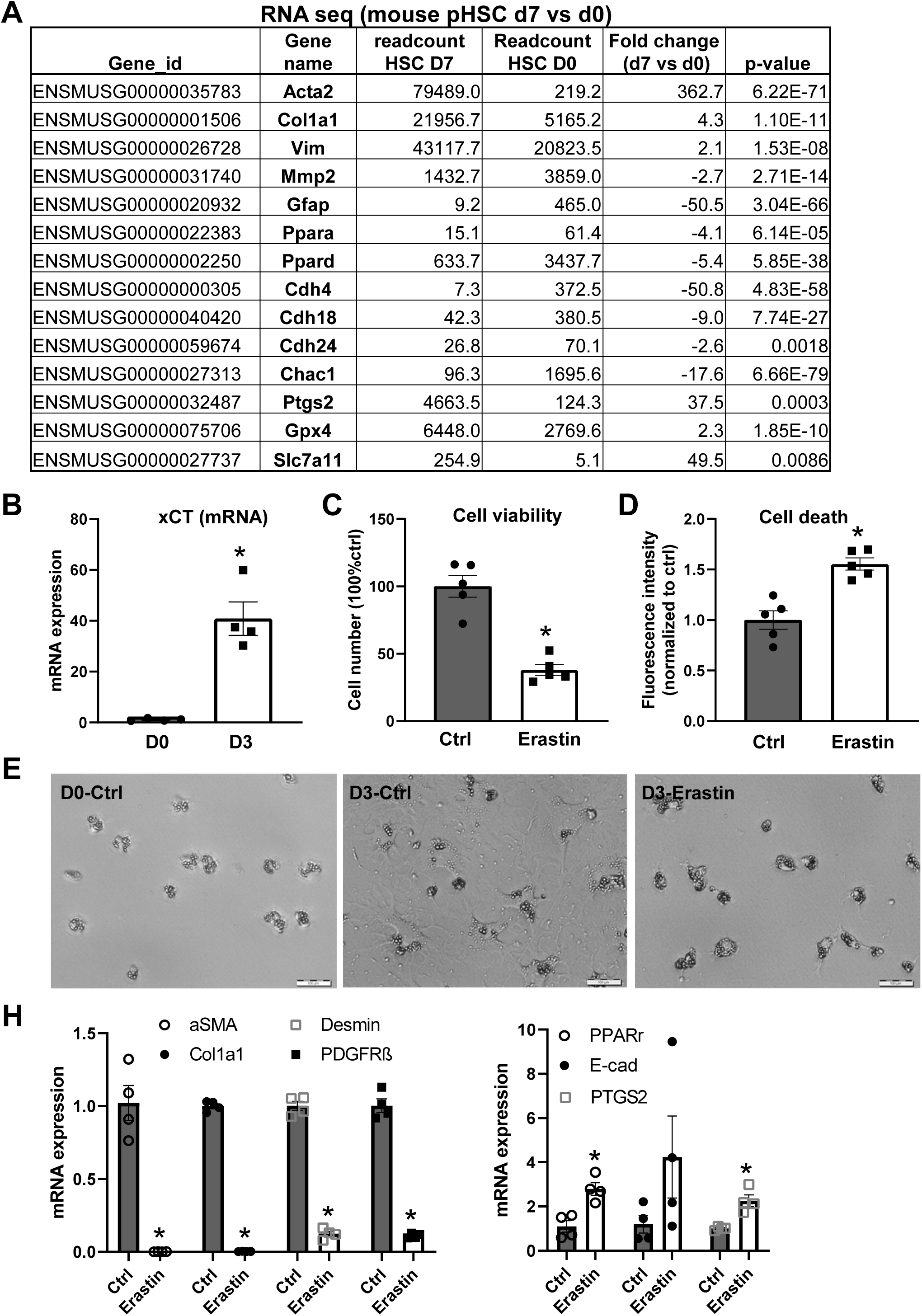

**Suppl. Fig. 11.**
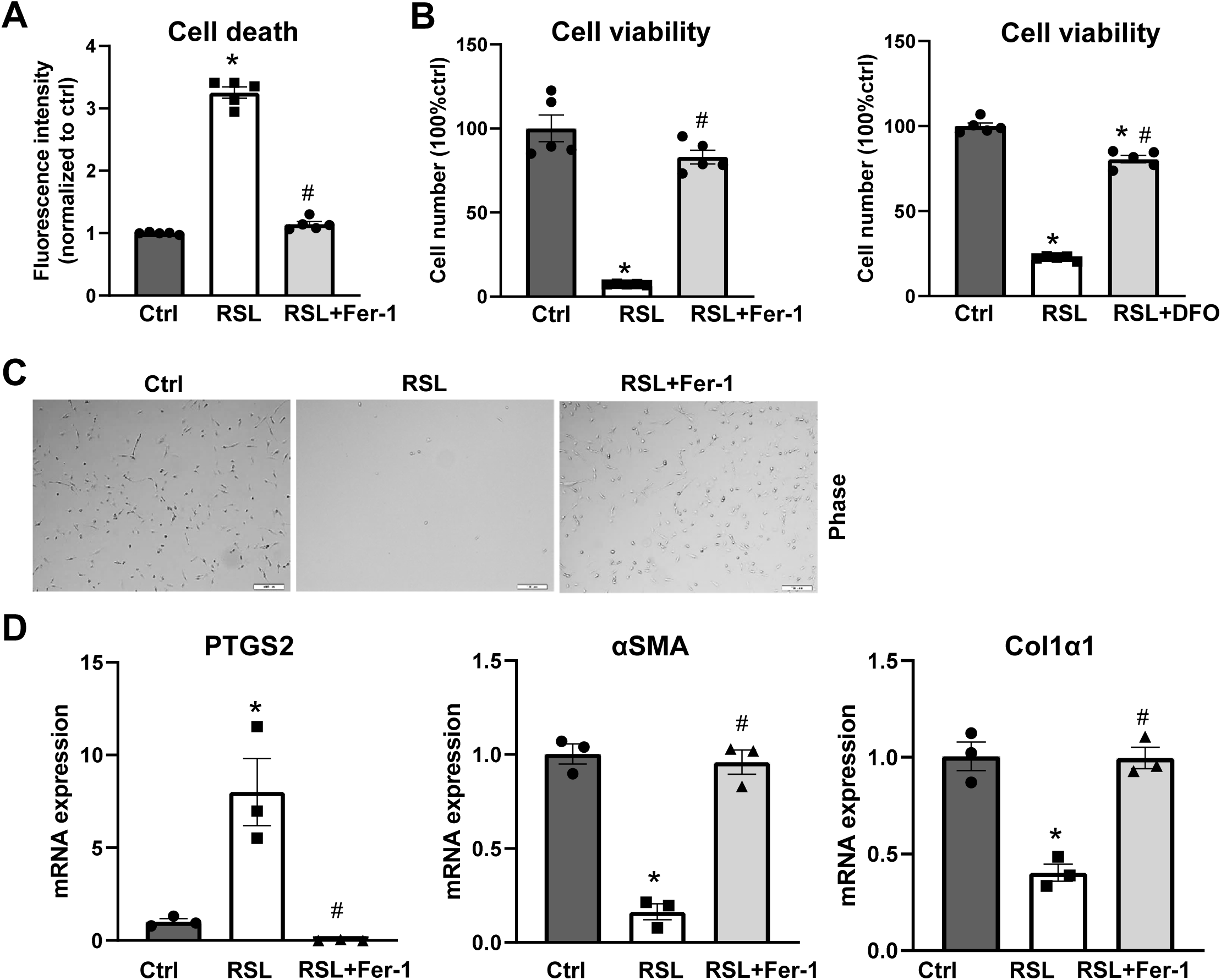

**Suppl. Fig. 12.**
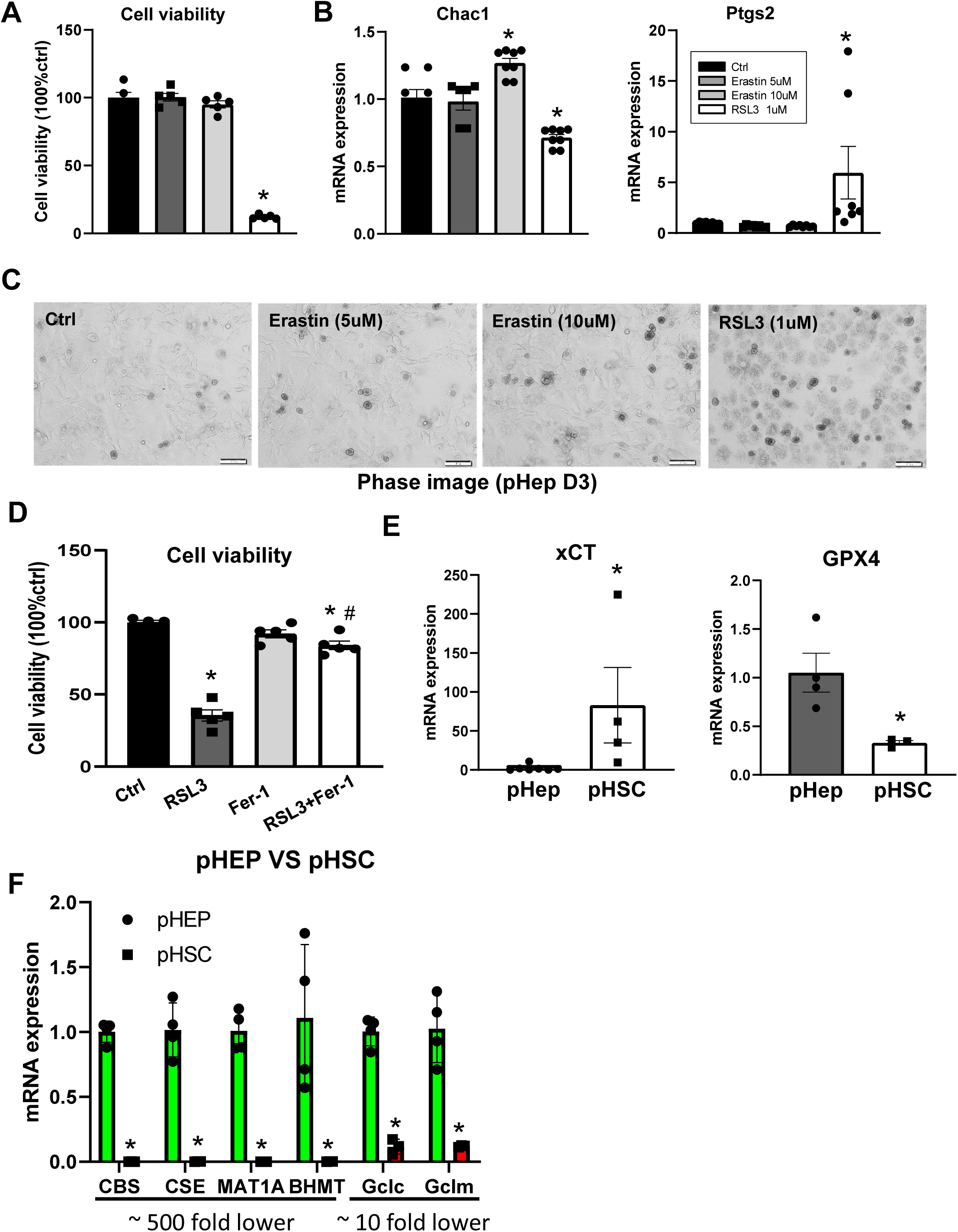

**Suppl. Fig. 13.**
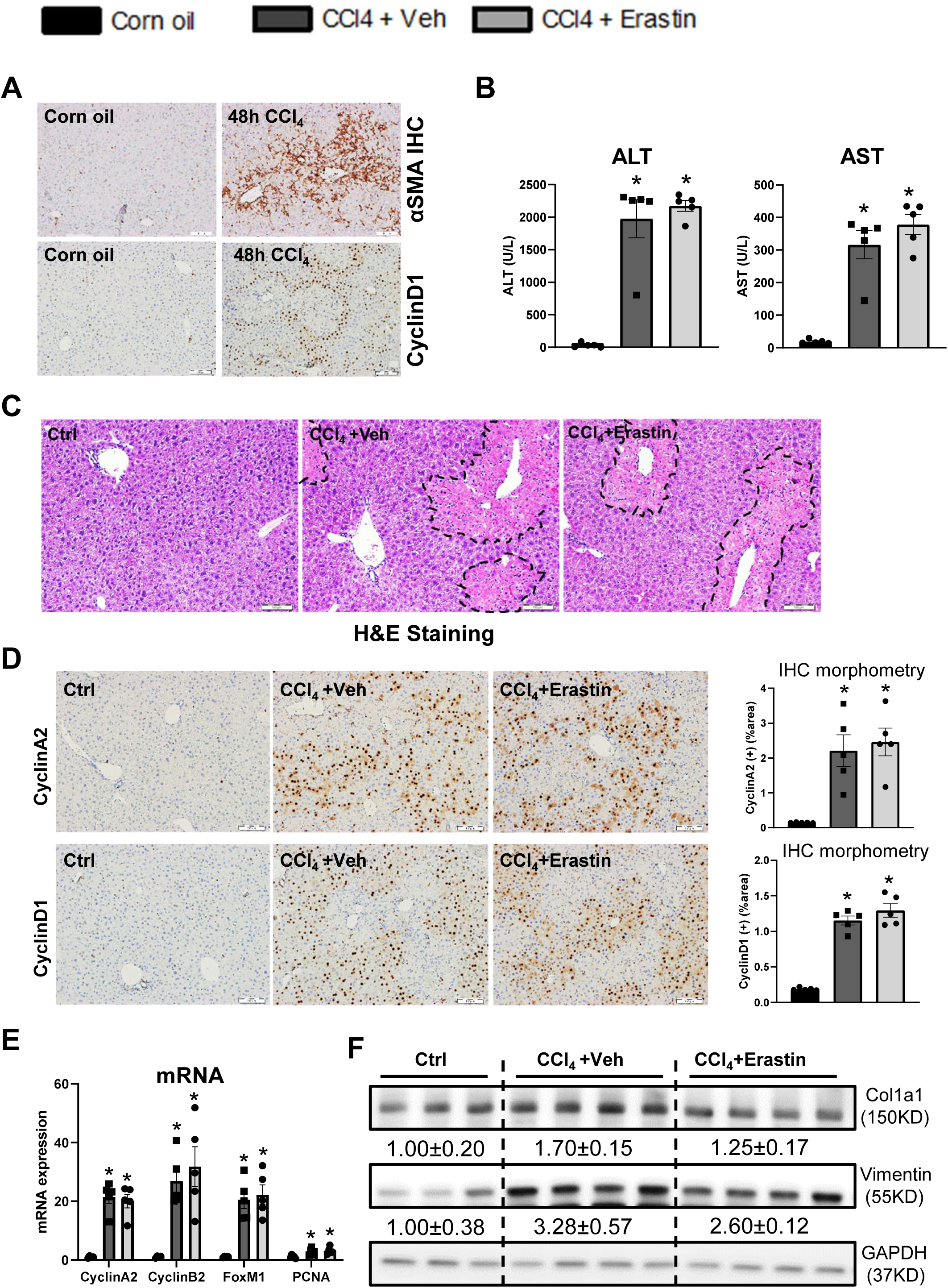

