## Supplemental materials for "Inhibiting xCT/SLC7A11 induces ferroptosis of myofibroblastic hepatic stellate cells and protects against liver fibrosis"

**Supplemental Figure 1. xCT deficiency suppressed growth and fibrogenic activity of MF-HSCs.** Human MF-HSCs (LX2 cells) cells were treated with 20nM non-targeting siRNA or xCT siRNA for 5 days. **(A)** Representative phase pictures at day 5. **(B)** mRNA of PPARγ and E-cad was quantified by qRT-PCR. Bars represent mean ± SEM of n = 3 assays. *$p < 0.05$ vs non-targeting RNA group.

**Supplemental Figure 2. Pharmacologically inhibiting xCT suppressed growth and fibrogenic activity of MF-HSCs.** Human MF-HSCs (LX2 cells) or rat MF-HSCs (8B cells) were treated with xCT inhibitor erastin, SASP or vehicle ctr (0.1% DMSO) for up to 4 days. **(A)** Representative phase pictures of LX2 cells at day 4. **(B)** Cell growth was determined by CCK8 assay. **(C)** Representative phase pictures of 8B cells treated with 250 μM SASP, 1 μM erastin or their vehicle ctrl for 4 days. **(D)** mRNA was quantified by qRT-PCR at day 4 in rat 8B cells treated with 1 μM erastin. **(E)** Protein expression of αSMA and Col1 was determined by ICC in rat 8B cells treated with 1 μM erastin. Bars represent mean ± SEM of n = 3-5 assays. *$p < 0.05$ vs ctr group.

**Supplemental Figure 3. Suppressed cell growth of MF-HSC caused by erastin was rescued by antioxidants.** 8B MF-HSC cells were treated 1 μM erastin or vehicle (0.1% DMSO) for 3 days; in some experiments, the culture medium was supplemented with 2 mM NAC or 50 μM β-ME. **(A)** Cell growth was determined by CCK8 assay. **(B)** Representative phase pictures of LX2 cells. Bars represent mean ± SEM of n = 5 assays. *$p < 0.05$ vs ctr group. $^{\#}p < 0.05$ vs erastin alone group.

**Supplemental Figure 4. Suppressed myofibroblastic activity of MF-HSC caused by SASP was rescued by antioxidants.** 8B MF-HSC cells were treated 250 μM SASP or vehicle (0.1% DMSO) for 3 days; in some experiments, the culture medium was supplemented with 2 mM NAC or 50 μM β-ME. **(A)** Cell growth was determined by CCK8 assay. **(B)** Representative phase pictures of LX2 cells. **(C)** Protein expression of αSMA and Vim was determined by western blotting. **(D-F)** mRNA of αSMA, Col1α1 and PPARγ was quantified by qRT-PCR. **(G)** Protein expression of Col1 was determined by ICC. Bars represent mean ± SEM of n = 4-5 assays. *$p < 0.05$ vs ctr group. $^{\#}p < 0.05$ vs SASP alone group.

**Supplemental Figure 5. xCT inhibition induced MF-HSC ferroptosis. (A)** Human MF-HSCs (LX2 cells) cells were treated with 20nM non-targeting siRNA (si-NT) or xCT siRNA (si-xCT). At day 5, xCT mRNA was quantified by qRT-PCR. Bars represent mean ± SEM of n = 3 assays. *p < 0.05 vs si-NT group. **(B-C)** LX2 cells were treated with SASP (500 μM) or vehicle ctr (0.1% DMSO), while 8B cells were treated with erastin (1 μM) or SASP (250 μM) or vehicle ctr (0.1% DMSO); some cells were supplemented with the canonical ferroptosis inhibitor ferrostatin (2 μM), or apoptosis inhibitor Z-VAD-FMK (10 μM) or necroptosis inhibitor Necrostain-1 (0.5 μM). Lipid ROS was assessed by C11-BODIPY assay (B) and representative phase pictures of these cells were acquired. **(D)** LX2 cells were treated with xCT inhibitor erastin (2 μM) or vehicle ctr (0.1% DMSO) for 4 days; some cells were supplemented with antioxidant NAC (2 mM) or β-ME (50 μM), caspase 3 activation was assessed by western blotting; **(E)** mRNA of CHAC1 was quantified by qRT-PCR. Bars represent mean ± SEM of n = 3 assays. *p < 0.05 vs ctrl group. $^{\#}$p < 0.05 vs erastin alone group.

**Supplemental Figure 6. xCT inhibition suppressed myofibroblastic HSC phenotype before causing cell death.** Human MF-HSCs (LX2 cells) cells were treated with xCT inhibitor erastin (2 μM) or SASP (500 μM) or vehicle ctr (0.1% DMSO) for 18h. **(A)** Cell death was assessed using CellTox™ Green Cytotoxicity assay. Cell lysis buffer was included as the positive control for complete cell death. **(B-D)** mRNA of αSMA, Col1α1 and MMP2 was quantified by qRT-PCR. Bars represent mean ± SEM of n = 4-5 assays. *p < 0.05 vs ctrl group.

**Supplemental Figure 7. MAPK pathway is involved in erastin-induced ferroptosis MF-HSCs.** MF-HSCs (LX2 or 8B cells) were treated with xCT inhibitor erastin or vehicle ctr (0.1% DMSO) for up to 4 days; some cells were supplemented with the JNK inhibitor SP600125 (10 μM) and p38 kinase inhibitor SB202190 (10 μM). **(A, D)** Cell growth was determined by CCK8 assay. **(B)** Cell death was assessed using CellTox™ Green Cytotoxicity assay. Cell lysis buffer was included as the positive control for complete cell death. **(C, E)** Representative phase pictures at day 4. Bars represent mean ± SEM of n = 5 assays. *p < 0.05 vs ctr group. $^{\#}$p < 0.05 vs erastin group.

**Supplemental Figure 8. Glutaminolysis derived glutamate is critical for xCT-mediated cystine uptake in MF-HSCs**. LX2 cells were treated with GLS1 inhibitor CB839 (1μM) with or without the supplement of 2 mM NAC or 50 μM β-ME for up to 4 days. **(A)** GSH levels were determined at 2h after treatment. **(B)** Cell growth was determined by CCK8 assay. **(C)** mRNA of αSMA was quantified by qRT-PCR at day 4. **(D)** Representative phase pictures of LX2 cells at day 4. 8B cells were treated with GLS1 inhibitor BPTES (10 μM) with or without the supplement of 2 mM NAC or 50 μM β-ME for up to 4 days. **(E)** Cell growth was determined by CCK8 assay. **(F)** Representative phase pictures at day 4. In following experiments, MF-HSCs (LX2 or 8B cells) were treated with BPTES (10 μM) or xCT inhibitor erastin or both for up to 4 days. **(G)** Cell death was assessed using CellTox™ Green Cytotoxicity assay. Cell lysis buffer was included as the positive control to induce complete cell death. **(H, J)** Cell growth was determined by CCK8 assay and corresponding representative phase pictures of at day 4. **(I, K)** mRNA of αSMA and Col1α1 was quantified by qRT-PCR at day 4. Bars represent mean ± SEM of n = 4-5 assays. *p < 0.05 vs CB839 or BPTES alone group.

**Supplemental Figure 9. Inhibiting xCT decreases GLS1 expression, while inhibiting GLS1 increases xCT expression**. **(A, C)** LX2 cells were treated with 20nM non-targeting siRNA, or xCT siRNA, or GLS1 siRNA for 5 days. **(B)** LX2 cells were treated with xCT inhibitor erastin (2 μM) or SASP (500 μM) or vehicle ctr (0.1% DMSO) for 3 days. **(D)** LX2 cells were treated with GLS1 inhibitor CB839 (1μM) or BPTES (10 μM) or vehicle ctr (0.1% DMSO) for 3 days. mRNA of xCT and GLS1 was quantified by qRT-PCR. Bars represent mean ± SEM of n = 3-4 assays. *p < 0.05 vs non-targeting RNA group or vehicle ctrl group.

**Supplemental Figure 10. Primary HSCs are sensitive to xCT inhibition. (A)** Primary HSCs were isolated from healthy C57Bl/6 male mice. A portion of the pooled isolate was harvested for Q-HSCs (Day 0; D0) and remaining cells were cultured for 7 days to induce trans-differentiation. RNA was isolated and RNA expression was compared at D7 versus D0 by RNA sequencing. **(B)** Primary HSCs were isolated from adult mice and then cultured for 3 days. Cells were harvested for xCT mRNA analysis using qRT-PCR. In following experiments, primary HSCs were treated with erastin (5 μM) for 3 days beginning immediately after isolation. **(C)** Cell viability was assessed by CCK8 assay. **(D)** Cell death was assessed using CellTox™ Green Cytotoxicity assay. **(E)** Representative phase pictures. **(H)** mRNA was quantified by qRT-PCR. Bars represent mean ± SEM of n=4-5 assays. *p < 0.05 vs D0 or vehicle ctrl at D3.

**Supplemental Figure 11. MF-HSCs are vulnerable to GPX4 inhibition.** Human MF-HSCs (LX2 cells) cells were treated with GPX inhibitor RSL3 (0.5 μM) or vehicle ctrl (0.1% DMSO) for 2 days. In some experiments, the culture medium was supplemented with the canonical ferroptosis inhibitor Fer-1 (2 μM), or iron chelator deferoxamine (10 μM). **(A)** Cell death was assessed using CellTox™ Green Cytotoxicity assay. **(B)** Cell viability was determined by CCK8 assay. **(C)** Representative phase pictures were acquired at day 2. **(D)** mRNA of PTGS2, αSMA and Col1 was quantified by qRT-PCR. Bars represent mean ± SEM of n = 3-5 assays. *$p < 0.05$ vs ctrl group. $^{\#}p < 0.05$ vs RSL3 group.

**Supplemental Figure 12. Primary hepatocytes are resistant to xCT inhibition, but vulnerable to GPX4 inhibition.** Primary mouse hepatocytes were treated with xCT inhibitor (5 μM; 10 μM) or GPX inhibitor RSL3 (1 μM) or vehicle ctrl (0.1% DMSO) for 3 days. In some experiments, the culture medium was supplemented with the canonical ferroptosis inhibitor Fer-1 (2 μM)**.** **(A)** Cell viability was determined by CCK8 assay. **(B)** mRNA of Chac1 and Ptgs2 was quantified by qRT-PCR. **(C)** Representative phase pictures of the cells at day 3. **(D)** Cell viability was determined by CCK8 assay. **(E, F)** Relative mRNA expression of indicated gene normalized to housekeeping gene S9 in primary HSCs compared to hepatocytes. Bars represent mean ± SEM of n = 4-8 assays. *$p < 0.05$ vs ctrl group (panel A, B, D), or vs pHep (panel E). $^{\#}p < 0.05$ vs RSL3 group.

**Supplemental Figure 13. xCT inhibition did not affect hepatocyte injury or regeneration.** Adult mice were injected intraperitoneally with corn oil or $CCl_4$. At 9h and 30h post-$CCl_4$ injection, some mice were intraperitoneally injected with erastin or its vehicle (10% DMSO), and liver tissues were harvested at 48h post-$CCl_4$. **(A)** Representative αSMA and CyclinD1 immuno-stained liver sections harvested from mice 48 h after treatment with either vehicle (Corn oil) or $CCl_4$. **(B)** Serum alanine aminotransferase (ALT) and aspartate aminotransferase (AST) levels 48 h after treatment with vehicle or CCl4. **(C)** Representative H&E-stained liver sections with injured areas outlined. Injured areas were identified by lack of nuclear staining or pyknotic nuclei and lines were marked along the boundaries of injured areas of parenchyma. **(D)** Representative cyclinA2 and cyclinD1 stained liver sections and corresponding morphometric analysis. **(E)** mRNA levels of proliferative marker cyclinA2, cyclinE1, FoxM1 and PCNA determined by qRT-PCR. **(F)** Protein expression of Col1α1 and vimentin as quantified by western blotting with GAPDH as the loading control. Bars represent mean ± SEM of n=5 mice/group. *$p < 0.05$ vs corn oil.

**Supplementary Table 1.** Primer used for qRT-PCR

| | Gene symbol | Primer Forward | Primer Reverse |
|---|---|---|---|
| **Human** | *S9* | GACTCCGGAACAAACGTGAGGT | CTTCATCTTGCCCTCGTCCA |
| | *PPARγ* | CGTGGCCGCAGATTTGAA | CTTCCATTACGGAGAGATCCAC |
| | *Acta2 (αSma)* | GGAGATCACGGCCCTAGCAC | AGGCCCGGCTTCATCGTAT |
| | *Col1a1 (Col1α1)* | CGGTGTGACTCGTGCAGC | ACAGCCGCTTCACCTACAGC |
| | *Desmin* | CAGTGGCTACCAGGACAACA | GCTGGTTTCTCGGAAGTTGA |
| | *E-Cadherin* | TGCCCAGAAAATGAAAAAGG | GTGTATGTGGCAATGCGTTC |
| | *MMP2* | CTGGGAGCATGGCGATGGATA | GGAAGCGGAATGGAAACTTG |
| | *CHAC1* | CCTGAAGTACCTGAATGTGCGAGA | GCAGCAAGTATTCAAGGTTGTGGC |
| | *PTGS2* | ATATGTTCTCCTGCCTACTGGAA | GCCCTTCACGTTATTGCAGATG |
| | *xCT (SLC7A11)* | ATGCAGTGGCAGTGACCTTT | GGCAACAAAGATCGGAACTG |

| | | | |
|---|---|---|---|
| **Mouse** | *S9* | GGGCCTGAAGATTGAGGATT | CGGGCATGGTGAATAGATTT |
| | *Chac1* | CATAGGGGCAGCGACAAGATG | CTGTGTGGCAATGACCTCTTC |
| | *Ptgs2* | CTGCGCCTTTTCAAGGATGG | GGGGATACACCTCTCCACCA |
| | *xCT* | TGCAATCAAGCTCGTGAC | AGCTGTATAACTCCAGGGACTA |
| | *Col1a1 (Col1α1)* | GAGCGGAGAGTACTGGATCG | GCTTCTTTTCCTTGGGGTTC |
| | *Col3a1 (Col3α1)* | GGTTTCTTCTCACCCTTCTTC | CTTCCAGACATCTCTAGACTCA |
| | *Col6a1 (Col1α1)* | CGACTGCGCCATTAAGAAG | CGGTCACCACGATCAAGTA |
| | *Acta2 (αSma)* | GCCAGTCGCTGTCAGGAACCC | AGCCGGCCTTACAGAGCCCA |
| | *Desmin* | TACACCTGCGAGATTGATGC | ACATCCAAGGCCATCTTCAC |
| | *Vim (Vimentin)* | GCCGAAAGCACCCTGCAGTCA | TGGGCCTGCAGCTCCTGGAT |

| | | | |
|---|---|---|---|
| **Rat** | *S9* | *GACTCCGGAACAAACGTGAGGT* | *CTTCATCTTGCCCTCGTCCA* |
| | *Chac1* | *GCCCTGTGGATTTTCGGGTA* | *ATCTTGTCGCTGCCCCTATG* |
| | *Ptgs2* | *GATTGACAGCCCACCAACTT* | *ACGTGGGGAGGGTAGATCAT* |
| | *MMP2* | *TGTGTCTTCCCCTTCACTTTTCTG* | *CGGTCATCATCGTAGTTGGTTGTG* |
| | *Pparg (Pparγ)* | *CCCTGGCAAAGCATTTGTAT* | *ACTGGCACCCTTGAAAAATG* |
| | *Vim (Vimentin)* | *GCCGAGGAATGGTACAAGT* | *CTCTTCCATTTCACGCATCT* |
| | *Acta2 (αSma)* | *GTGGGGGACGAAGCGCAGAG* | *GGCCTTAGGGTTCAGCGGCG* |
| | *Col1a1 (Col1α1)* | *CTGCATACACAATGGCCTAA* | *GGGTCCCTCGACTCCTA* |
